# STAMBPL1 activates the GRHL3/HIF1A/VEGFA axis through interaction with FOXO1 to promote angiogenesis in triple-negative breast cancer

**DOI:** 10.1101/2024.08.31.610659

**Authors:** Huan Fang, Huichun Liang, Chuanyu Yang, Dewei Jiang, Qianmei Luo, Wenming Cao, Huifeng Zhang, Ceshi Chen

## Abstract

In the clinic, anti-tumor angiogenesis is commonly employed for treating recurrent, metastatic, drug-resistant triple-negative and advanced breast cancer. Our previous research revealed that the deubiquitinase STAMBPL1 enhances the stability of MKP-1, thereby promoting cisplatin resistance in breast cancer. In this study, we discovered that STAMBPL1 could upregulate the expression of the hypoxia-inducible factor HIF1α in breast cancer cells. Therefore, we investigated whether STAMBPL1 promotes tumor angiogenesis. We demonstrated that STAMBPL1 increased *HIF1A* transcription in a non-enzymatic manner, thereby activating the HIF1α/VEGFA signaling pathway to facilitate TNBC angiogenesis. Through RNA-seq analysis, we identified the transcription factor GRHL3 as a downstream target of STAMBPL1 that is responsible for mediating *HIF1A* transcription. Furthermore, we discovered that STAMBPL1 regulates *GRHL3* transcription by interacting with the transcription factor FOXO1. These findings shed light on the role and mechanism of STAMBPL1 in the pathogenesis of breast cancer, offering novel targets and avenues for the treatment of triple-negative and advanced breast cancer.

## Introduction

Triple-negative breast cancer (TNBC) is a subtype of breast cancer characterized by a lack of expression of estrogen receptor (ER), progesterone receptor (PR), and human epidermal growth factor receptor 2 (HER2). Despite representing only 10–15% of all breast cancer cases, it is associated with a greater risk of recurrence, metastasis, and resistance to chemotherapy, leading to a poorer prognosis than other types of breast cancer[1]. Therefore, understanding the pathogenesis of this subtype of breast cancer and identifying effective treatment targets are key areas of research focus and challenge. In the clinic, in addition to surgery, radiotherapy, and chemotherapy, treatment approaches for TNBC also include the use of epidermal growth factor receptor (EGFR) inhibitors, poly (ADP-ribose) polymerase (PARP) inhibitors, immune checkpoint inhibitors, and anti-angiogenic therapies[2, 3].

For solid tumors, the survival, proliferation, and invasion of cancer cells rely on surrounding blood vessels to provide nutrients and oxygen. Cancer cells can increase the synthesis and secretion of the vascular endothelial growth factor A (VEGFA), activate endothelial cells in neighboring blood vessels through paracrine pathways, and stimulate tumor angiogenesis[4]. Therefore, investigating the upstream molecular mechanisms that activate VEGFA in cancer cells offers new targets for inhibiting TNBC angiogenesis via disruption of this signaling pathway. The deubiquitinase STAMBPL1 (also known as AMSH-LP) belongs to the AMSH family and cleaves K63-linked polyubiquitin chains[5]. STAMBPL1 stabilizes XIAP to inhibit apoptosis in prostate cancer cells[6] and confers resistance to honokiol-induced apoptosis in various cancer cell types by stabilizing Survivin and c-FLIP[7]. Our previous research demonstrated that STAMBPL1 enhances cisplatin resistance in TNBC cells through the stabilization of MKP-1[8].

In this study, we discovered that, independent of its deubiquitinating enzyme activity, STAMBPL1 upregulates HIF1α expression in TNBC cells. We aimed to investigate the molecular mechanism by which it upregulates HIF1α and explore its role in tumor angiogenesis in TNBC.

FOXO1, a member of the forkhead box transcription factor family, regulates various physiological and pathological processes, such as cell proliferation, apoptosis, autophagy, and oxidative stress[9]. In breast cancer, FOXO1 enhances the stemness of cancer cells by promoting *SOX2* transcription[10] and confers resistance to chemotherapy drugs in basal-like breast cancer cells by activating *KLF5* transcription[11]. Furthermore, FOXO1 plays a crucial role in angiogenesis by upregulating VEGFA transcription[12]. However, the specific mechanism by which FOXO1 regulates VEGFA transcription remains unclear, and its role in tumor angiogenesis in TNBC requires further investigation.

In this study, we discovered that STAMBPL1 facilitates the transcriptional regulation of *GRHL3* by interacting with FOXO1, consequently enhancing *HIF1A* transcription through GRHL3 to activate the HIF1α/VEGFA pathway. This results in increased endothelial cell activity via paracrine signaling, thereby promoting tumor angiogenesis in TNBC.

## Methods

### Cell lines and reagents

All cell lines used in this study, including HCC1806, HCC1937, and HEK293T cells, were purchased from ATCC (American Type Culture Collection, Manassas, VA, USA) and validated by short tandem repeat (STR) analysis, and these cell lines tested negative for mycoplasma contamination. HCC1806 and HCC1937 cells were cultured in RPMI 1640 medium supplemented with 5% FBS. HEK293T cells were cultured in DMEM (Thermo Fisher, Grand Island, USA) with 5% FBS at 37°C with 5% CO_2_. Primary human umbilical vein endothelial cells (HUVECs) were maintained in an EGM-2 Bullet Kit (CC-3162, Lonza, USA). AS1842856 (Cat#HY-100596) and apatinib (Cat#HY-13342S) were purchased from MCE (New Jersey, USA).

### Plasmid construction and stable STAMBPL1, GRHL3 and FOXO1 overexpression

We constructed the full-length *STAMBPL1/GRHL3/FOXO1* genes and then subcloned them into the pCDH lentiviral vector. The packaging plasmids (pMDLg/pRRE, pRSV-Rev, and pCMV-VSV-G) and the pCDH-STAMBPL1/GRHL3/FOXO1 expression plasmid were cotransfected into HEK293T cells (2 ×10^6^ in 10 cm plates) to produce lentiviruses. Following transfection for 48 hours, the lentivirus was collected and used to infect HCC1806 and HCC1937 cells. Forty-eight hours later, puromycin (2 μg/ml) was used to screen the cell populations.

### Stable knockdown of STAMBPL1 and GRHL3

The pSIH1-H1-puro shRNA vector was used to express STAMBPL1, GRHL3 and luciferase (LUC) shRNAs. *STAMBPL1* shRNA#1, 5’-GGAGCATCAGAGATTGATA-3’; *STAMBPL1* shRNA#2, 5’-GCTGCTATGCCTGACCATA-3’; *GRHL3*shRNA#1, 5’-CCTTGAGCTTCCTCTATGA-3’; *GRHL3* shRNA#2, 5’-AGAGGAAGATGCGCGATGA-3’; *Luciferase* shRNA, 5’-CUUACGCUGAGUACUUCGA-3’; HCC1806 and HCC1937 cells were infected with lentivirus. Stable populations were selected via the use of 1 to 2 mg/mL puromycin. The knockdown effect was evaluated by Western blotting.

### RNA interference

The siRNA target sequences used in this study were as follows: *STAMBPL1* siRNA#1, 5’-GGAGCATCAGAGATTGATA-3’; *STAMBPL1* siRNA#2, 5’-GCTGCTATGCCTGACCATA-3’; *GRHL3* siRNA#1, 5’-CCTTGAGCTTCCTCTATGA-3’; *GRHL3* siRNA#2, 5’-AGAGGAAGATGCGCGATGA-3’; *HIF1α* siRNA#1, 5’-AAGAGGTGGATATGTCTGG-3’; *HIF1α* siRNA#2, 5’-CGTCGAAAAGAAAAGTCTCTT-3’; *FOXO1*siRNA#1, 5’-CCCAGAUGCCUAUACAAAC-3’; and *FOXO1* siRNA#2, 5’-CTCAAATGCTAGTACTATTAG-3’. All siRNAs were synthesized by RiboBio (RiboBio, China) and transfected at a final concentration of 50 nM.

### Western blotting (WB) and antibodies

The WB procedure was described in our previous study[13]. Anti-STAMBPL1 (sc-376526) and anti-GAPDH (sc-25778) antibodies were purchased from Santa Cruz Biotechnology (Santa Cruz, CA, USA). Anti-FOXO1 (#2880S) and anti-HIF1α (#36169S) antibodies were purchased from CST. Anti-β-actin (A5441) and anti-GST (G7781) antibodies were purchased from Sigma‒ Aldrich (St. Louis, MO, USA). The anti-Flag (ab205606) antibody was purchased from Abcam.

### Real-time polymerase chain reaction (RT‒qPCR)

Total RNA was extracted via TRIzol (Invitrogen, 15596026). One microgram of total RNA was reverse transcribed to cDNA according to the manufacturer’s instructions for the HiScript II QRT SuperMix for qPCR Kit (Vazyme, R223-01). For quantitative PCR (RT‒qPCR), the SYBR Green Select Master Mix system (Applied Biosystems, 4472908, USA) was used on an ABI-7900HT system (Applied Biosystems, 4351405). The primers used for PCR were as follows: 18S forward, 5′-CTCAACACGGGAAACCTCAC-3′; 18S reverse, 5′-CGCTCCACCAACTAAGAACG-3′; HIF1α forward, 5′-AAGTCTGCAACATGGAAGGTAT-3′; HIF1α reverse, 5′-TGAGGAATGGGTTCACAAATC-3′; VEGFA forward, 5′-TGAGGAATGGGTTCACAAATC-3′; VEGFA reverse, 5′-ATCTGCATGGTGATGTTGGA-3′; GLUT1 forward, 5′-TCGTCGGCATCCTCATCGCC-3′; GLUT1 reverse, 5′-CCGGTTCTCCTCGTTGCGGT-3′; GRHL3 forward, 5′-GGGGCTGAGGAATGCGATCT-3′; and GRHL3 reverse, 5′-AATTTTGCCGTCCAGCTCCC-3′.

### Cell proliferation and migration assays

To detect the proliferation of HUVECs, we used the Click-iT™ EdU Alexa Fluor™ 647 Imaging Kit (Invitrogen) according to the manufacturer’s protocol. Briefly, HUVECs were seeded on coverslips (BD Biosciences) at 0.5×10^5^ cells per well. The next day, the supernatants were discarded, and the cells were cultured with conditioned medium (CM). Six hours later, the cells were incubated with EdU in CM for 4 hours, followed by fixation and staining. For each sample, three random fields were observed via fluorescence microscopy, and the total numbers of cells and EdU-positive cells were counted. To detect the migration of HUVECs, we performed a wound-healing assay. Twenty-four hours after seeding, the supernatants of the HUVECs were discarded, and the cells were scratched and cultured with CM for 24 hours. Wound closure was imaged via microscopy. For each image, the gap width was analyzed via ImageJ.

### Tube formation assays

HUVECs (1 × 10^4^) in CM were seeded onto Matrigel (BD Biosciences)-coated μ-Slide angiogenesis plates (ibidiGmbH, Munich, Germany). At 6 hours after seeding, images were taken via microscopy and then analyzed with ImageJ. The total branch length was measured.

### Chromatin immunoprecipitation assays

After the sample preparation was completed, the subsequent experimental steps were performed according to the instructions of the Simple ChIP (R) Plus Kit (CST, # 9005). The PCR primers for amplifying the region of interest on the HIF1α gene promoter were as follows: 5′-GACTGACAGGCTTGAAGTTTATGC-3′ and 5′-TGTTGCTGTAAACTTCAAGGGAAA-3′, and the PCR primers for amplifying the region of interest on the GRHL3 gene promoter were as follows: 5′-TTCTATCCCTTCTGTGCTGACCA-3′ and 5′-TGTGCCAGACCCTACTCTGGG-3′.

### Dual-luciferase reporter assays

The DNA fragments HIF1α and GRHL3 were amplified from MCF10A cell genomic DNA via PCR template and then cloned and inserted into the pGL3-Basic vector. HEK293T cells were seeded in 24-well plates and transfected with pCDH-GRHL3-3×Flag or pCDH-FOXO1-3×Flag and pGL3 luciferase reporter plasmids (both 600 ng/well) together with the pCMV-Renilla control (5 ng/well). After transfection for 48 h, the cell lysates were collected, and the luciferase activities were detected via the dual-luciferase reporter assay system (Promega, USA). For the WT HIF1α promoter (with the GRHL3 binding motif), the primers used for PCR were as follows: forward, 5′-gctagcccgggctcgagatctCCACTGCGCTCCAGCCTG-3′; reverse, 5′-cagtaccggaatgccaagcttCCTCAGACGAGGCAGCACTG-3′. For the mutant HIF1α promoter (without the GRHL3 binding motif), the primers used for PCR were as follows: forward, 5′-TCTTTCCCTGAGGCCTTCCTATATGCTTAT-3′; reverse, 5′-ATAAGCATATAGGAAGGCCTCAGGGAAAGA-3′. For the WT GRHL3 promoter (with the FOXO1 binding motif), the primers used for PCR were as follows: forward, 5′-gctagcccgggctcgagatctATTAACAAGGGTGACTGAAGAGGG-3′; reverse, 5′-cagtaccggaatgccaagcttTGGAGGTATACCTCAACAGGTGC-3′. For the mutant GRHL3 promoter (without the FOXO1 binding motif), the primers used for PCR were as follows: forward, 5′-CTCCCCCACCAAACAAAGAAGGAGAACACCCC-3′; reverse, 5′-GGGGTGTTCTCCTTCTTTGTTTGGTGGGGGAG-3′.

### Immunofluorescence staining

HEK293T cells plated on cell culture slides were transfected with the STAMBPL1 and FOXO1 expression plasmids. Two days after transfection, the cells were fixed in 3.7% polyformaldehyde at 4°C overnight. After being blocked with 5% BSA at room temperature for 1 hour, the cells were stained with anti-STAMBPL1 (mouse) and anti-FOXO1 (rabbit) antibodies at 4°C overnight. The cells were subsequently stained with both an Alexa Fluor647-labeled anti-mouse secondary antibody and a FITC-labeled anti-rabbit secondary antibody (ZSGB-Bio, Beijing, China) at room temperature for 1 hour. Nuclei were stained with DAPI (Biosharp, BL739A) for 15 min. Images were captured via a confocal microscope.

### Immunoprecipitation and GST pull-down

For endogenous protein interaction, cell lysates were first incubated with anti-FOXO1 antibodies or rabbit IgG (2729S; Cell Signaling Technology) and then incubated with protein A/G magnetic beads (HY-K0202; MCE). For the GST pull-down assay, the cell lysates were directly incubated with Glutathione Sepharose 4B (10312185; Cytiva) overnight at 4°C. For the IP-Flag assay, cell lysates were directly incubated with anti-Flag magnetic beads (HY-K0207; MCE) overnight at 4°C. The precipitates were washed four times with 1 ml of lysis buffer, boiled for 10 minutes with 1×SDS sample buffer, and subjected to WB analysis.

### Xenograft experiments

We purchased 5-to 6-week-old female BALB/c nude mice from SLACCAS (Changsha, China). Animal feeding and experiments were approved by the animal ethics committee of Kunming Institute of Zoology, Chinese Academy of Sciences. HCC1806-shLuc, HCC1806-shSTAMBPL1, or HCC1806-shGRHL3 cells and HCC1806-PCDH-Vector or HCC1806-PCDH-STAMBPL1 cells (1 × 10^6^ in Matrigel (BD Biosciences, NY, USA)) were implanted into the mammary fat pads of the mice. When the tumor volume reached approximately 50 mm^3^, the nude mice were randomly assigned to the control or treatment groups (n = 4/group). The control group was given vehicle alone, and the treatment group received the FOXO1 inhibitor AS1842856 (10 mg/kg), the VEGFR inhibitor apatinib (50 mg/kg), and the FOXO1 inhibitor AS1842856 combined with the VEGFR inhibitor apatinib via intragastric administration every two days for 20 days. The tumor volume was calculated as follows: tumor volume was calculated by the formula (π × length × width^2^)/6. The maximal tumor size permitted by the animal ethics committee of Kunming Institute of Zoology, Chinese Academy of Sciences, and the maximal tumor size in this study was not exceeded.

### Immunohistochemical staining

The xenograft tumor tissues were fixed in 3.7% formalin solution. The immunohistochemistry was performed on 4-μm-thick paraffin sections after pressure-cooking for antigen retrieval. An anti-CD31 primary antibody (1:400, Abcam, ab28364) was used. After 12 h, the slides were washed three times with PBS and incubated with secondary antibodies (hypersensitive enzyme-labeled goat anti-mouse/rabbit IgG polymer (OriGene, China) at room temperature for 20 min, DAB concentrated chromogenic solution (1:200 dilution of concentrated DAB chromogenic solution)), counterstained with 0.5% hematoxylin, dehydrated with graded concentrations of ethanol for 3 min each (70%–80%–90%–100%), and finally stained with dimethyl benzene. The immunostained slides were evaluated via light microscopy, and the number of microvessels with positive CD31 expression was counted.

### Statistical analysis

All the graphs were created via GraphPad Prism software version 8.0. Comparisons between two independent groups were assessed via two-tailed Student’s *t* tests. One-way analysis of variance with least significant differences was used for multiple group comparisons. *P* values of < 0.05, 0.01 or 0.001 were considered to indicate statistically significant results, and comparisons that were significant at the 0.05 level are indicated by *, those at the 0.01 level are indicated by **, and those at the 0.001 level are indicated by ***.

## Results

### STAMBPL1 upregulates HIF1α in a non-enzymatic manner in TNBC cells

Our previous research revealed that STAMBPL1, a deubiquitinase, promotes cisplatin resistance in TNBC by stabilizing MKP1[8]. To further investigate the role and mechanism of STAMBPL1 in TNBC, we discovered that the knockdown of STAMBPL1 can inhibit hypoxia-induced HIF1α expression (Fig.1A) and suppress the transcription of the HIF1α downstream genes *VEGFA* and *GLUT1* (Fig.1B-E). Under normoxic conditions, both STAMBPL1 and its enzymatically mutated variant (E292A) can upregulate the protein expression of HIF1α (Fig.1F). These findings suggest that STAMBPL1 enhances the expression of HIF1α in breast cancer cells through a non-DUB enzyme activity mechanism.

**Figure 1.**
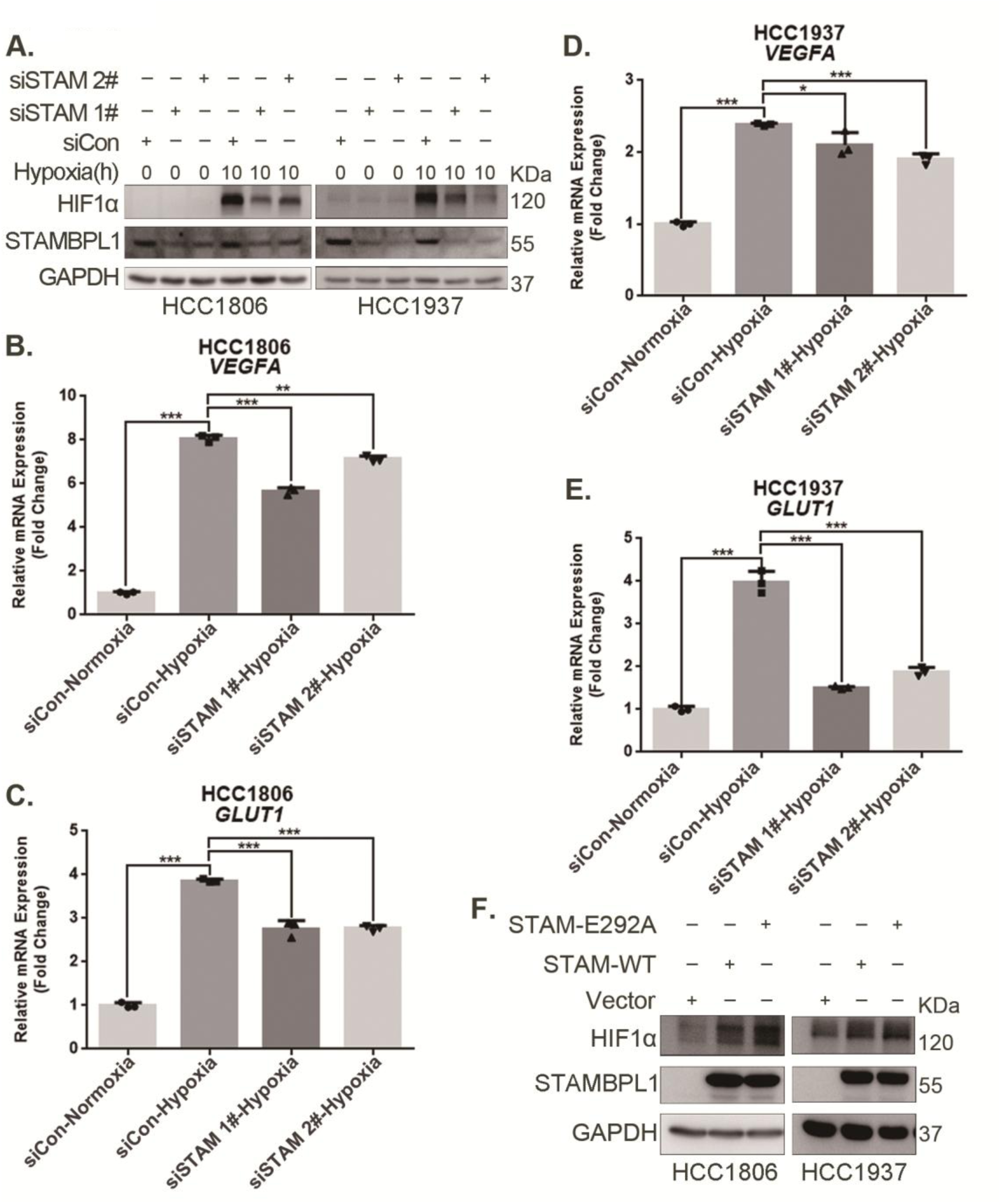
STAMBPL1 upregulates HIF1α in a non-enzymatic manner in TNBC cells. **(A)** Western blot detected the protein levels of HIF1α when STAMBPL1 was knocked down under normoxia and hypoxia for 10 hours. **(B-C)** RNA samples were collected after 10 h of hypoxia treatment following the knockdown of STAMBPL1 in HCC1806 cells. RT‒qPCR experiments were performed to detect the mRNA levels of the HIF1α downstream targets *VEGFA* and *GLUT1*. **(D-E)** RNA samples of HCC1937 were collected as the above HCC1806 cells. RT‒ qPCR experiments were performed to detect the mRNA levels of the HIF1α downstream targets *VEGFA* and *GLUT1*. **(F)** Western blot detected the protein levels of HIF1α when STAMBPL1 /STAMBPL1-E292A was overexpressed under normoxia in HCC1806 and HCC1937 cells. *: *P*<0.05, **: *P*<0.01, ***: *P*<0.001, *t* test.

### STAMBPL1 promotes *HIF1A* transcription and activates the HIF1α/VEGFA axis

To elucidate how STAMBPL1 upregulates HIF1α, we examined the transcription levels of *HIF1A* when STAMBPL1 was knocking down under hypoxia. These results indicate that knocking down STAMBPL1 significantly inhibits the transcription of *HIF1A* (Fig.2A-C). At the same time, we detected the transcription level of HIF1A and VEGFA when STAMBPL1 was overexpressing under normoxia. Data showed that STAMBPL1 overexpression increased the transcription of *HIF1A* (Fig.4E and Fig. S3E) and *VEGFA* (Fig.2E and 2G). The ability of STAMBPL1 to induce *VEGFA* transcription was blocked by HIF1α knockdown (Fig.2D-G). These findings suggested that STAMBPL1 activated the HIF1α/VEGFA axis through enhancing the transcription of *HIF1α*.

**Figure 2.**
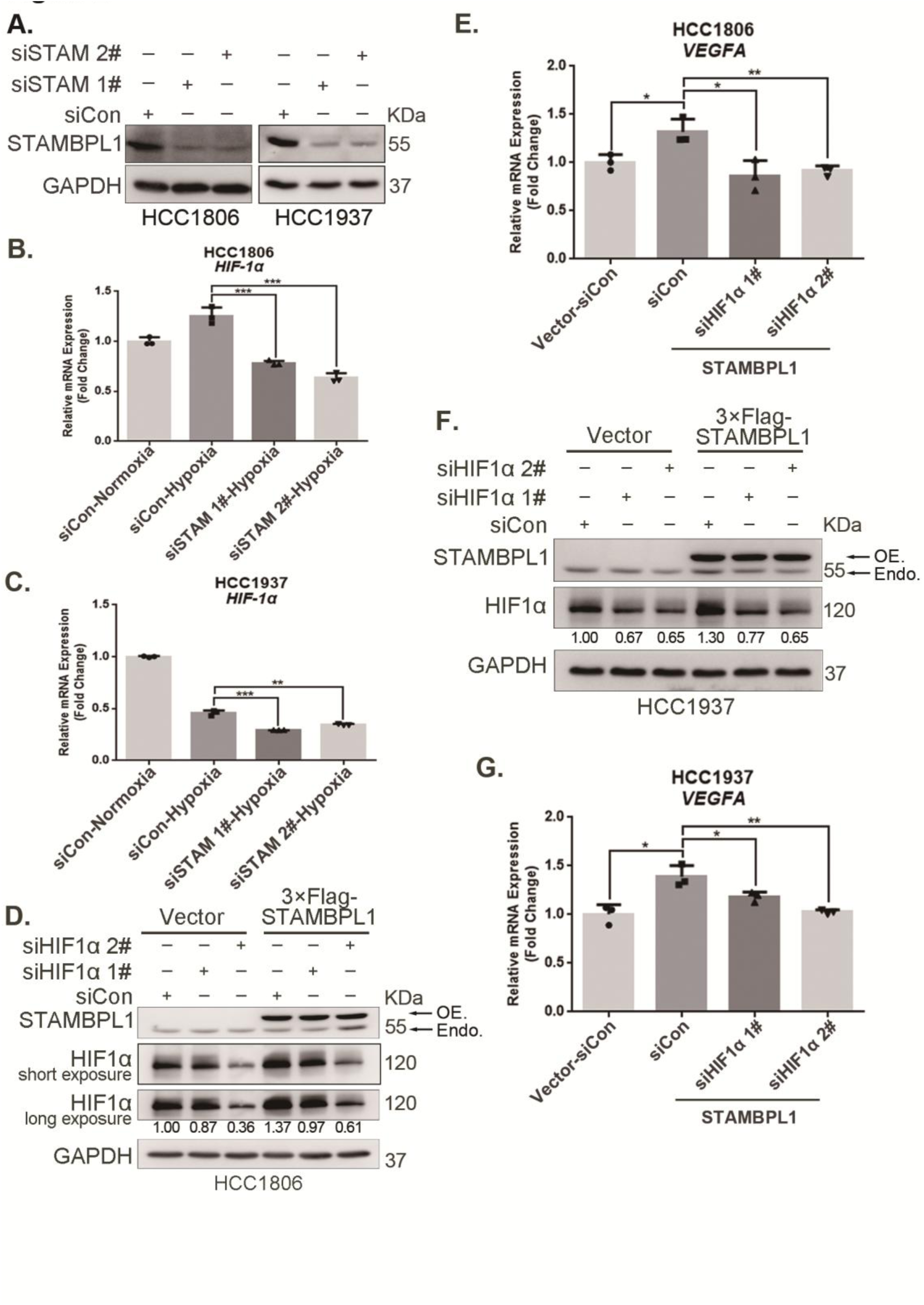
STAMBPL1 promotes *HIF1A* transcription and activates the HIF1α/VEGFA axis. **(A)** Western blot analysis was performed to confirm the knockdown of STAMBPL1 in HCC1806 and HCC1937 cells. **(B)** RNA samples were collected after 10 h of hypoxia treatment following the knockdown of STAMBPL1 in HCC1806 cells. RT‒qPCR experiments were performed to detect the mRNA levels of HIF1α. **(C)** RNA samples of HCC1937 cells were collected as the above HCC1806 cells. RT‒qPCR experiments were performed to detect the mRNA levels of HIF1α. **(D)** Knocking down HIF1α using siRNAs in HCC1806 cells stably overexpressing STAMBPL1 under normoxia. Western blot was used to detect the protein levels of STAMBPL1 and HIF1α. **(E)** RT‒qPCR experiments were performed to detect the effect of HIF1α knockdown on the mRNA level of *VEGFA* mRNA in HCC1806 cells with STAMBPL1 overexpression under normoxia. **(F)** Knocking down HIF1α using siRNAs in HCC1937 cells stably overexpressing STAMBPL1 under normoxia. Western blot was used to detect the protein levels of STAMBPL1 and HIF1α. **(G)** RT‒qPCR experiments were performed to detect the effect of HIF1α knockdown on the mRNA level of *VEGFA* mRNA in HCC1937 cells with STAMBPL1 overexpression under normoxia. *: *P*<0.05, **: *P*<0.01, ***: *P*<0.001, *t-*test.

### STAMBPL1 in TNBC cells enhances the activity of HUVECs and promotes TNBC angiogenesis

The conditioned medium (CM) from TNBC cells with STAMBPL1 knockdown inhibited the proliferation (Fig.3A-B, and Fig.S1A-B), migration (Fig.3C-D, and Fig.S1C-D), and tube formation (Fig.3E-F, and Fig.S1E-F) of HUVECs. When STAMBPL1 was overexpressed in TNBC cells, the conditioned medium of HCC1806 and HCC1937 cells promoted the ability of HUVECs to proliferate (Fig.S2A and Fig.S2D), migrate (Fig.S2B and Fig.S2E), and form tubes (Fig.S2C and Fig.S2F), which could be reversed by knocking down HIF1α in TNBC cells. These findings suggest that STAMBPL1 activates the HIF1α/VEGFA axis in TNBC cells, leading to enhanced abilities of HUVECs through a paracrine pathway. Knocking down STAMBPL1 inhibited the growth of HCC1806 xenografts in mice (Fig.3G-I) and decreased the number of microvessel in tumor tissue (Fig.3J-K), indicating that STAMBPL1 promoted tumor angiogenesis.

**Figure 3.**
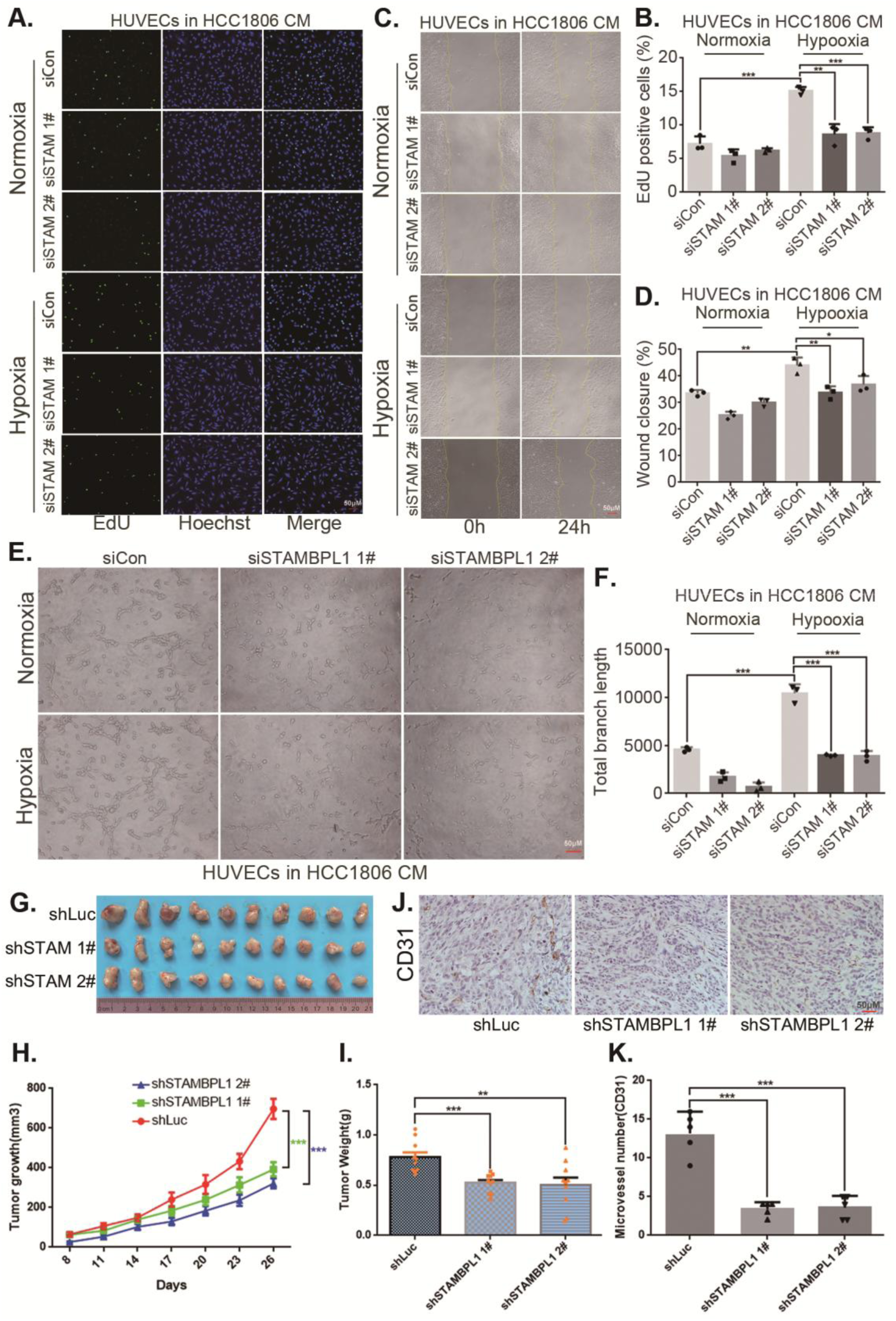
STAMBPL1 in TNBC cells enhances the activity of HUVECs and promotes TNBC angiogenesis. **(A)** EDU assay was performed to detect the effect of HCC1806 conditioned medium (CM) with STAMBPL1 knockdown under normoxia or hypoxia on the proliferation of HUVECs. **(B)** Statistical analysis of the EdU assay results. **(C)** Wound healing assay was performed to detect the effect of HCC1806 CM with STAMBPL1 knockdown under normoxia or hypoxia on the migration of HUVECs. **(D)** Statistical analysis of the wound healing assay results. **(E)** Tube formation assay was performed to detect the effect of HCC1806 CM with STAMBPL1 knockdown under normoxia or hypoxia on the tube formation of HUVECs. **(F)** Statistical analysis of the tube formation assay results. **(G)** The growth of TNBC xenograft tumors was evaluated by photographing the tumors in nude mice to assess the role of STAMBPL1. **(H-I)** The growth and weight of the transplanted tumors in the nude mice were statistically analyzed, and the STAMBPL1sh#1 and STAMBPL1sh#2 groups were compared with the shLuc group. **(J)** An immunohistochemical assay was used to detect the expression of the angiogenesis marker CD31 in xenograft tumors. **(K)** The number of microvessels in the immunohistochemical experiments was statistically analyzed. *: *P*<0.05, **: *P*<0.01, ***: *P*<0.001, *t-*test.

**Figure 4.**
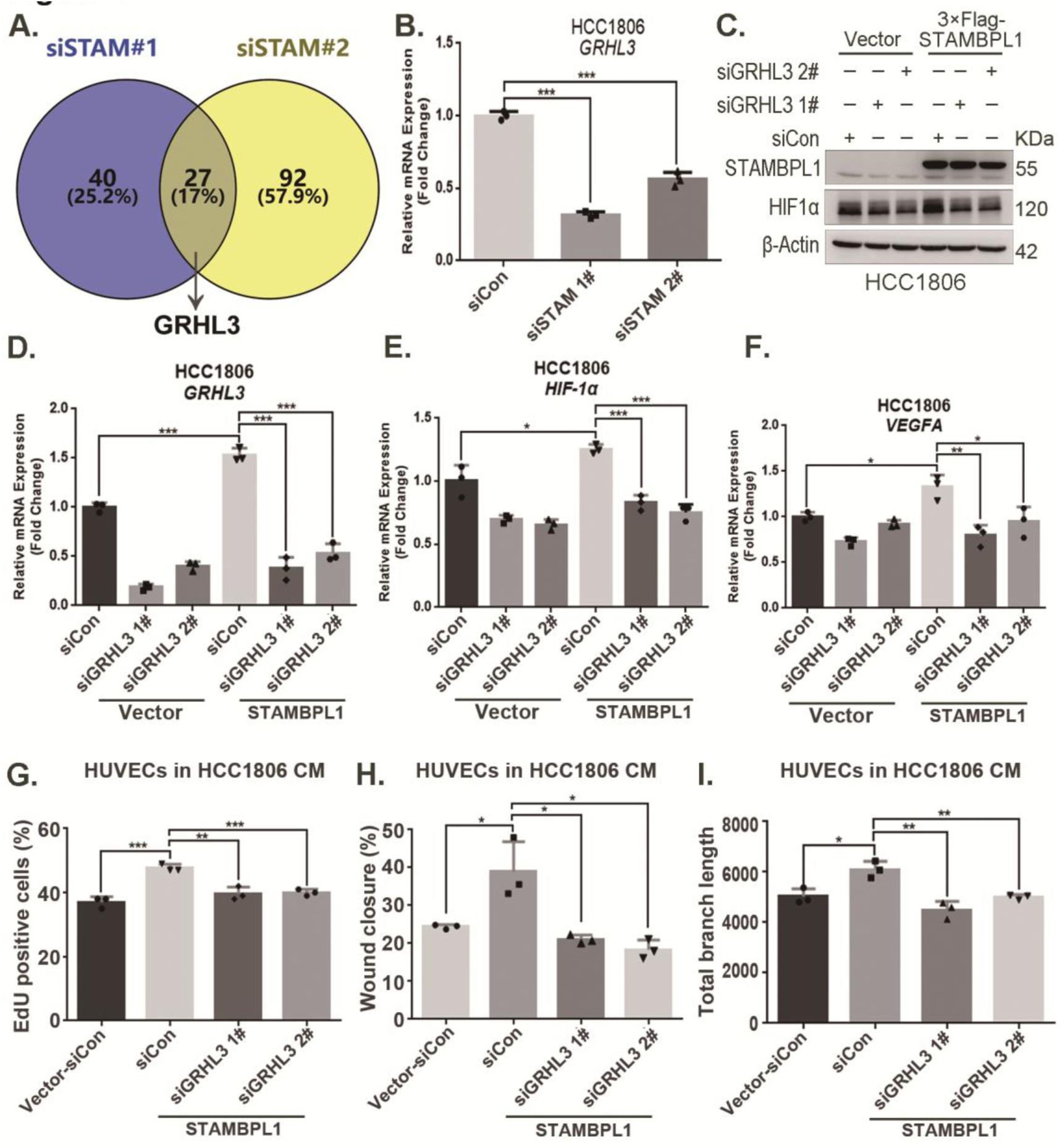
STAMBPL1 promotes *HIF1A* transcription via the upregulation of GRHL3. **(A)** Venn diagram showing the overlap of differentially expressed genes identified via RNA-seq analysis. **(B)** RT‒qPCR experiment was performed to detect the effect of STAMBPL1 knockdown on the mRNA levels of GRHL3 in HCC1806 cells. **(C)** In HCC1806 cells stably overexpressing STAMBPL1, GRHL3 was knocked down via siRNA, after which the protein was collected. Western blotting was performed to detect the effect of GRHL3 knockdown on the expression of HIF1α as STAMBPL1 overexpression under normoxia. **(D-F)** RT‒qPCR experiments were performed to detect the knockdown efficiency of GRHL3 and the effect of GRHL3 knockdown on the mRNA levels of HIF1α and its downstream target VEGFA when STAMBPL1 was overexpressing. **(G)** Statistical analysis of the EdU assay results. **(H)** Statistical analysis of the wound healing assay results. **(I)** Statistical analysis of the tube formation assay results. *: *P*<0.05, **: *P*<0.01, ***: *P*<0.001, *t-*test.

### STAMBPL1 promotes *HIF1A* transcription via the upregulation of GRHL3

By silencing the *STAMBPL1* gene in HCC1806 cells subjected to 10 hours of hypoxia, we performed RNA-seq analysis to investigate the mechanism by which STAMBPL1 promotes *HIF1A* transcription. We found that silencing of STAMBPL1 resulted in the downregulation of 27 genes (of which only 18 were annotated). Of these 18 genes, except for GRHL3, which is a transcription factor reported to be involved in gene transcription regulation, the remaining 17 genes have no documented association with RNA transcription, stability, or modification (Fig.4A and Fig.S3A). Silencing of STAMBPL1 inhibited *GRHL3* transcription in both HCC1806 and HCC1937 cells (Fig.4B and Fig.S3B). Conversely, the overexpression of STAMBPL1 promoted *GRHL3* transcription (Fig.4D and Fig.S3D). Furthermore, the stimulatory effects of STAMBPL1 on the HIF1α/VEGFA axis (Fig. 4C-F and Fig.S3C-F) and on HUVECs (Fig.4G-I and Fig.S3G-I) were reversed by GRHL3 knockdown. These findings suggest that STAMBPL1 activates the HIF1α/VEGFA axis by upregulating GRHL3.

### GRHL3 enhances *HIF1A* transcription by binding to its promoter

On the basis of the GRHL binding motif (Fig.5A), a GRHL binding sequence was identified in the *HIF1A* promoter (Fig.5B). ChIP and luciferase assays revealed that GRHL3 bound to the *HIF1A* promoter (Fig.5D) and increased its activity (Fig.5E). Knockdown of GRHL3 (Fig.5G and Fig.S4B) resulted in decreased expression of HIF1α at both the mRNA (Fig.5H and Fig.S4C) and protein levels (Fig.5F and Fig.S4A), leading to suppressed *VEGFA* transcription (Fig.5I and Fig.S4D), indicating that GRHL3 promotes *HIF1A* transcription by binding to its promoter.

**Figure 5.**
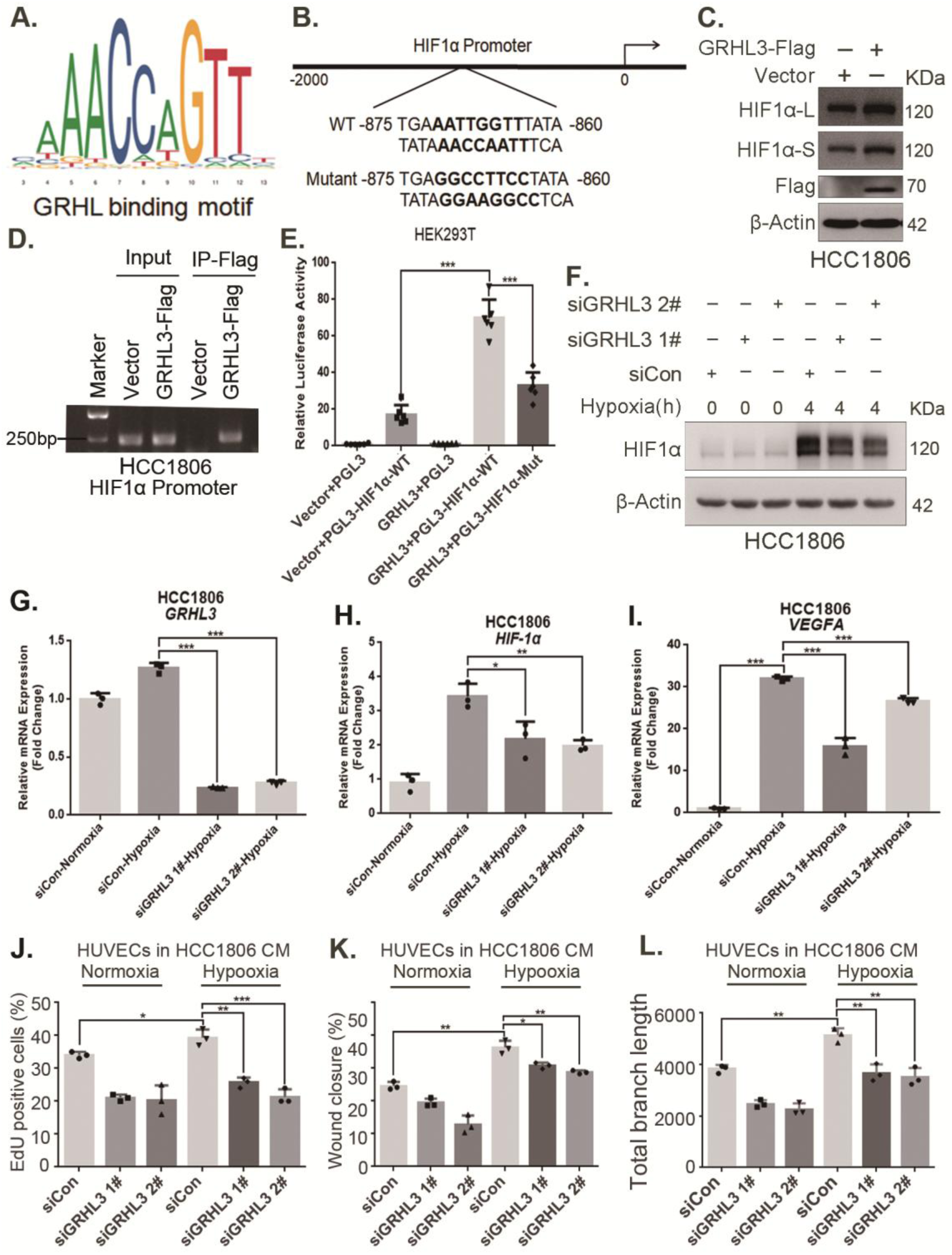
GRHL3 enhances *HIF1A* transcription by binding to its promoter. **(A)** The JASPAR website was used to predict the potential binding sequence of the transcription factor GRHL3 to the HIF1α promoter. **(B)** Mutation pattern of the HIF1α promoter binding sequence. **(C)** The protein level of HIF1α was detected in HCC1806 cells with GRHL3 overexpression by using Western blot. **(D)** ChIP‒PCR experiments conducted in HCC1806 cells stably overexpressing GRHL3 to detect the interaction between GRHL3 and the promoter of *HIF1A* gene. **(E)** A luciferase assay was performed in HEK293T cells to detect the effect of GRHL3 on the transcriptional activity of *HIF1A* promoter. **(F)** In HCC1806 cells, GRHL3 was knocked down by siRNA, and the cells were subjected to hypoxia for 4 hours. Western blotting was performed to detect the effect of GRHL3 knockdown on the protein level of HIF1α. **(G-I)** RT‒qPCR experiments were used to detect the effect of GRHL3 knockdown on the mRNA levels of HIF1α and its downstream target VEGFA. **(J)** In HCC1806 cells, GRHL3 was knocked down via siRNA, and the cells were then subjected to hypoxia for 24 hours. The conditioned medium was collected and used to treat HUVECs. EdU assays were performed to detect the effect of GRHL3 knockdown CM on the proliferation of HUVECs. Statistical analysis of the EdU assay results was performed. **(K)** Wound healing assays were performed to detect the effect of GRHL3 knockdown CM on the migration of HUVECs. Statistical analysis of the wound healing assay results was performed. **(L)** The tube formation assay was performed to detect the effect of GRHL3 knockdown CM on the tube formation of HUVECs. Statistical analysis of the tube formation assay results was performed. *: *P*<0.05, **: *P*<0.01, ***: *P*<0.001, *t-*test.

Furthermore, conditioned medium from TNBC cells with GRHL3 knockdown inhibited HUVEC proliferation, migration, and tube formation (Fig.5J-L and Fig.S4E-G). The overexpression of GRHL3 in TNBC cells activated the HIF1α/VEGFA axis (Fig.6A-B and Fig.S4H-J), resulting in increased proliferation, migration, and tube formation in HUVECs. This effect was reversed by knocking down HIF1α in TNBC cells (Fig.6C-E and Fig.S4K-M). Knockdown of GRHL3 (Fig.6F) also inhibited tumor growth in HCC1806 xenograft mice (Fig.6G-I) and reduced the number of microvessel in tumor tissues (Fig.6J-K).

**Figure 6.**
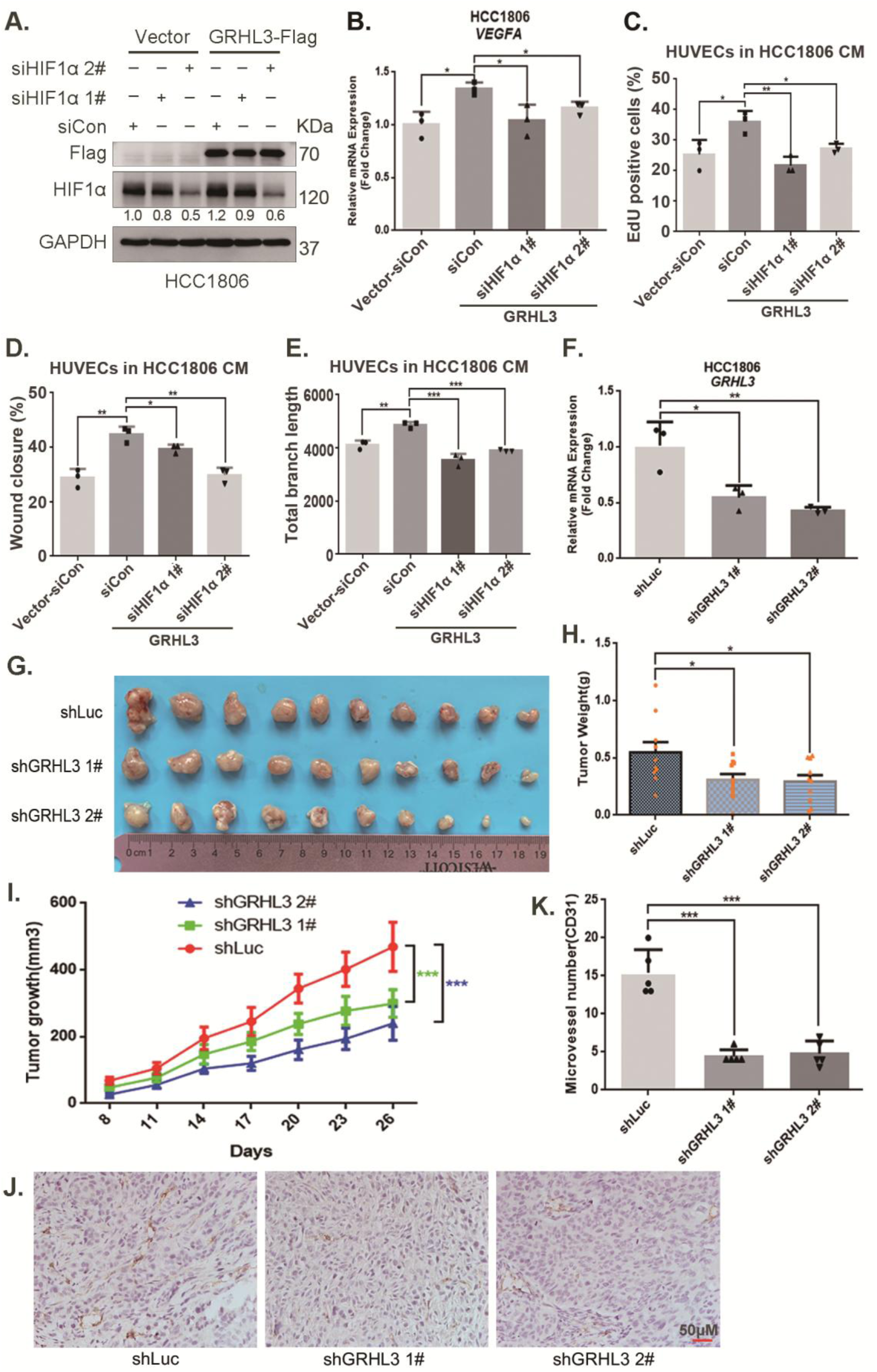
GRHL3 enhances *HIF1A* transcription by binding to its promoter. **(A)** In HCC1806 cells stably overexpressing GRHL3, HIF1α was knocked down by siRNA, and the protein was subsequently collected. Western blotting was performed to detect the protein level of GRHL3-Flag and HIF1α. **(B)** RT‒qPCR experiments were performed to detect the effect of HIF1α knockdown on the mRNA level of VEGFA as GRHL3 overexpression. **(C)** HCC1806 cells stably overexpressing GRHL3 were used for the knockdown of HIF1α via siRNA. The conditioned medium was then collected and used to treat HUVECs. EdU assay was performed to detect the effect of CM with HIF1α knockdown on the proliferation of HUVECs as GRHL3 overexpression. **(D)** Wound healing assay was performed to detect the effect of CM with HIF1α knockdown on the migration of HUVECs as GRHL3 overexpression. **(E)** The tube formation assay was performed to detect the effect of CM with HIF1α knockdown on the tube formation of HUVECs as GRHL3 overexpression. **(F)** RT‒qPCR was used to detect the knockdown of GRHL3 in HCC1806 cells. **(G)** Photographs of xenograft tumors from nude mice were taken to evaluate the role of GRHL3 in the growth of TNBC xenograft tumors. **(H)** The weights of the transplanted tumors from the nude mice were statistically analyzed, and the shGRHL3 1# and shGRHL3 2# groups were compared with the shLuc group. **(I)** The growth of transplanted tumors in nude mice was statistically analyzed, and the shGRHL3 1# and shGRHL3 2# groups were compared with the shLuc group. **(J)** An immunohistochemical assay was performed to detect the expression of the angiogenesis marker CD31 in xenograft tumors. **(K)** Statistical analysis of the number of microvessels in the immunohistochemical experiments. *: *P*<0.05, **: *P*<0.01, ***: *P*<0.001, *t-*test.

### STAMBPL1 mediates *GRHL3* transcription by interacting with FOXO1

Our previous study demonstrated that STAMBPL1 is localized in the nucleus[8]. This protein may promote the transcription of GRHL3 through interactions with other transcription factors. Previous studies have indicated that FOXO1 acts as an upstream transcription factor of GRHL3[14]. Therefore, we aimed to investigate whether STAMBPL1 promotes GRHL3 transcription via FOXO1. Our experimental data revealed that knockdown of FOXO1 in TNBC cells not only inhibited the GRHL3/HIF1α/VEGFA axis (Fig.7A-D and Fig.S5A-D) but also reversed the stimulatory effects of STAMBPL1 on this axis (Fig.7E-H). Furthermore, FOXO1 was found to bind to the promoter region of GRHL3 (Fig.7I and Fig.S5F) and enhance its transcriptional activity (Fig.7J). STAMBPL1 was shown to increase the transcriptional activation of FOXO1 at the GRHL3 promoter (Fig.7K), and it also bound to the GRHL3 promoter which could be disrupted by FOXO1 knockdown (Fig.7L). However, the overexpression of STAMBPL1 and its enzymatic activity mutants did not affect the protein expression level of FOXO1 (Fig.S5E). To investigate the mechanism by which STAMBPL1 promotes GRHL3 transcription through FOXO1, we conducted immunofluorescence and immunoprecipitation assays. We observed the colocalization of STAMBPL1 and FOXO1 in the nucleus (Fig.7M) and identified an interaction between them in HCC1937 cells (Fig.7N). Through the generation of a series of Flag-FOXO1/GST-fused STAMBPL1 deletion mutants, we mapped the regions of the proteins responsible for this interaction (Fig.S5G‒H). The results of the GST pulldown assay indicated that the N-terminus (1–140 aa) of STAMBPL1 interacted with FOXO1 (Fig.S5I). Additionally, an immunoprecipitation assay revealed an interaction between the FOXO1 protein (250–596 aa) and STAMBPL1 (Fig.S5J). These findings suggest that STAMBPL1 promotes GRHL3 transcription by interacting with FOXO1.

**Figure 7.**
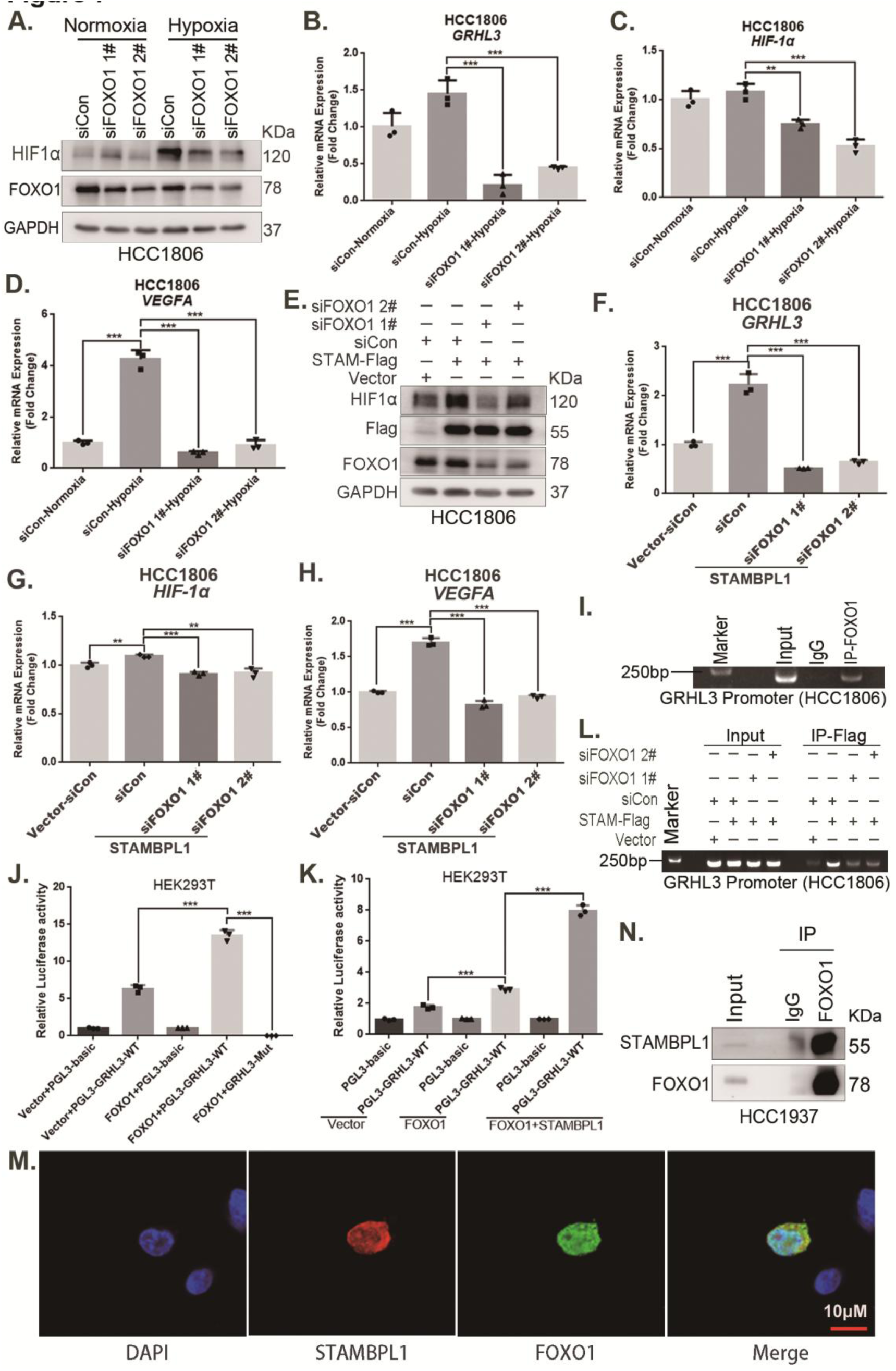
STAMBPL1 mediates *GRHL3* transcription by interacting with FOXO1. **(A)** Western blotting was used to detect the HIF1α protein expression when FOXO1 was knocking down in HCC1806 cells followed by hypoxia treatment for 4 hours. **(B-D)** RT‒qPCR experiments were performed to detect the effect of FOXO1 knockdown on the mRNA levels of GRHL3/HIF1α/VEGFA in HCC1806 cells under hypoxia for 4 hours. **(E)** In HCC1806 cells stably overexpressing STAMBPL1, protein samples were collected after FOXO1 was knocked down via siRNA. Western blotting was performed to detect the effect of FOXO1 knockdown on the expression of HIF1α as STAMBPL1 overexpression. **(F-H)** RT‒qPCR experiments were performed to detect the effect of FOXO1 knockdown on the mRNA expression of GRHL3, HIF1α and VEGFA as STAMBPL1 overexpression. **(I)** An endogenous ChIP‒PCR assay was performed using an anti-FOXO1 antibody in HCC1806 cells. **(J)** A luciferase assay was performed in HEK293T cells to detect the effect of FOXO1 on the transcriptional activity of GRHL3 promoter. **(K)** A luciferase assay was conducted in HEK293T cells to detected the effect of STAMBPL1 on the activation of the GRHL3 promoter by FOXO1. **(L)** ChIP‒PCR experiments were performed in HCC1806 cells stably overexpressing STAMBPL1 after knocking down FOXO1 via siRNA. **(M)** PCDH-STAMBPL1-3×Flag and PCDH-FOXO1-3×Flag plasmids were co-transfected into HEK293T cells, and then, immunofluorescence experiments were performed. Red represents STAMBPL1 staining, green represents FOXO1 staining, and blue represents DAPI staining. **(N)** Endogenous STAMBPL1 protein was immunoprecipitated from HCC1937 cells via an anti-FOXO1 antibody. Immunoglobulin G (IgG) served as the negative control. Endogenous STAMBPL1 was detected via western blotting. *: *P*<0.05, **: *P*<0.01, ***: *P*<0.001, *t-*test.

### The combination of VEGFR and FOXO1 inhibitors synergistically suppresses TNBC xenograft growth

Through an analysis of the Metabric database in BCIP (http://www.omicsnet.org/bcancer), it was observed that while the expression levels of STAMBPL1, FOXO1, and GRHL3 in breast cancer tissues are not universally elevated compared to adjacent non-cancerous tissues, but their expression levels in TNBC are significantly higher than those in non-TNBC (Fig.8A-C and Fig.S6A-C). To assess the potential therapeutic efficacy of cotreatment with the FOXO1 inhibitor AS1842856 and the VEGFR inhibitor apatinib *in vivo*, we conducted animal experiments in nude mice. HCC1806 cells overexpressing STAMBPL1 were orthotopically implanted into the mammary fat pads of 6-week-old female mice (n=16/group). Once the tumor volume reached approximately 50 mm^3^, the mice were divided into four subgroups to receive apatinib (50 mg/kg, once every two days), AS1842856 (10 mg/kg, once every two days), a combination of both drugs, or vehicle control for 20 days. Our findings indicated that STAMBPL1 overexpression enhanced breast cancer cell growth *in vivo*. While the individual inhibitory effects of the FOXO1 and VEGFR inhibitors on tumor growth were not significant, the combined treatment markedly suppressed tumor growth in nude mice (Fig.8E-G). Importantly, the drug treatments did not affect the body weights of the mice (Fig.8H). These results suggest that the combined administration of AS1842856 and apatinib effectively inhibits tumor growth in nude mice.

**Figure 8.**
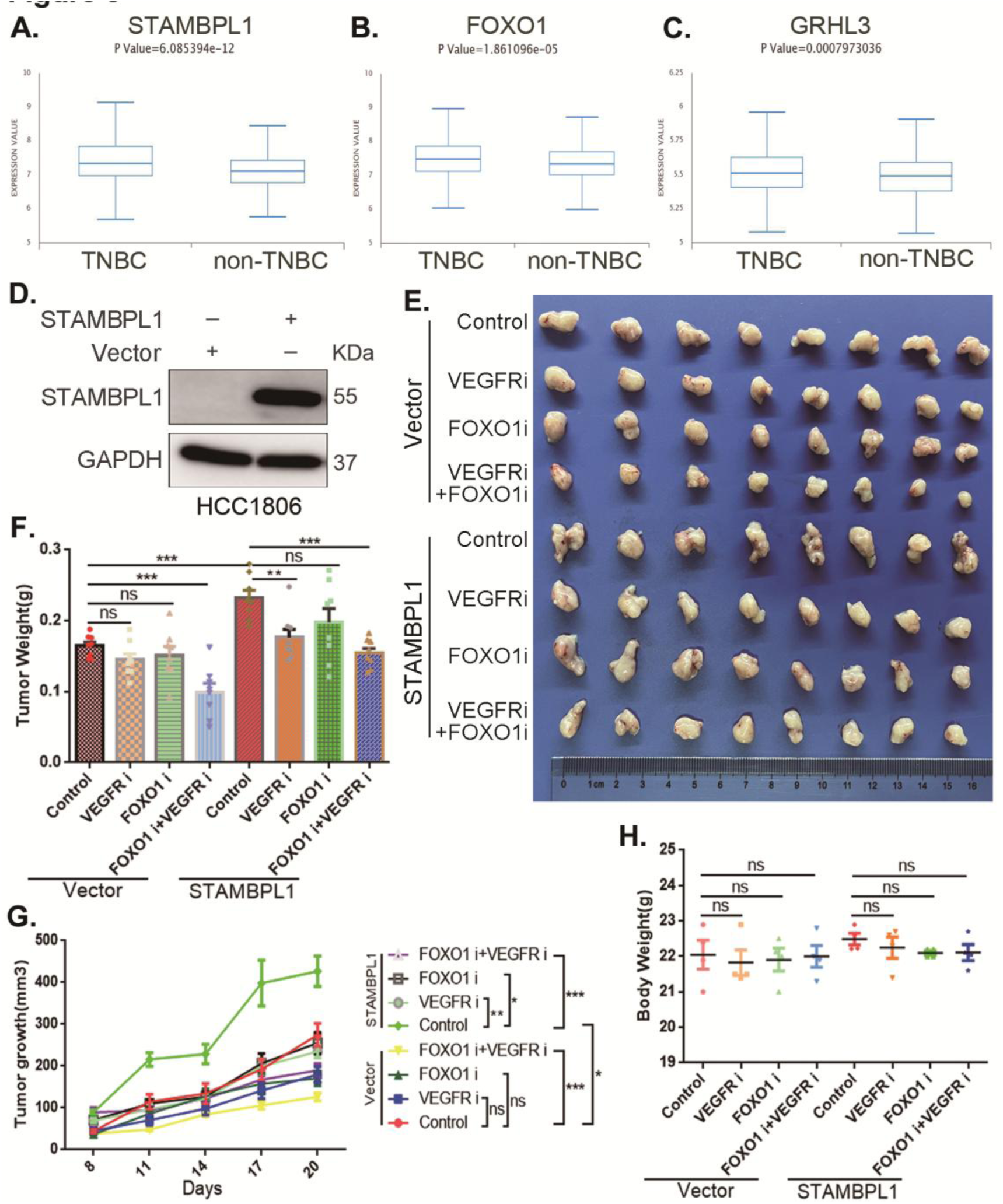
The combination of VEGFR and FOXO1 inhibitors synergistically suppresses TNBC xenograft growth. (**A-C**) Metabric database analysis in BCIP revealed high expression levels of STAMBPL1, FOXO1, and GRHL3 in TNBC (n = 319) compared with non-TNBC (n = 1661). **(D)** The effect of STAMBPL1 overexpression in HCC1806 cells was detected by Western blotting. **(E)** Nude mice with tumors received the FOXO1 inhibitor AS1842856 (10 mg/kg, every two days) or the VEGFR inhibitor Apatinib (50 mg/kg, every two days) or a combination of both treatments. The effect of the drug treatments on the transplanted tumors was assessed by imaging the tumors. **(F-G)** The weight and growth of the transplanted tumors in the nude mice were statistically analyzed. The vector control group and the STAMBPL overexpression group were compared. The nondrug group and the combined drug group in the vector control group were compared. The nondrug group and the combined drug group in the STAMBPL overexpression group were compared. **(H)** The final weights of the nude mice were statistically analyzed. *: *P*<0.05, **: *P*<0.01, ***: *P*<0.001, *t-*test.

**Figure 9.**
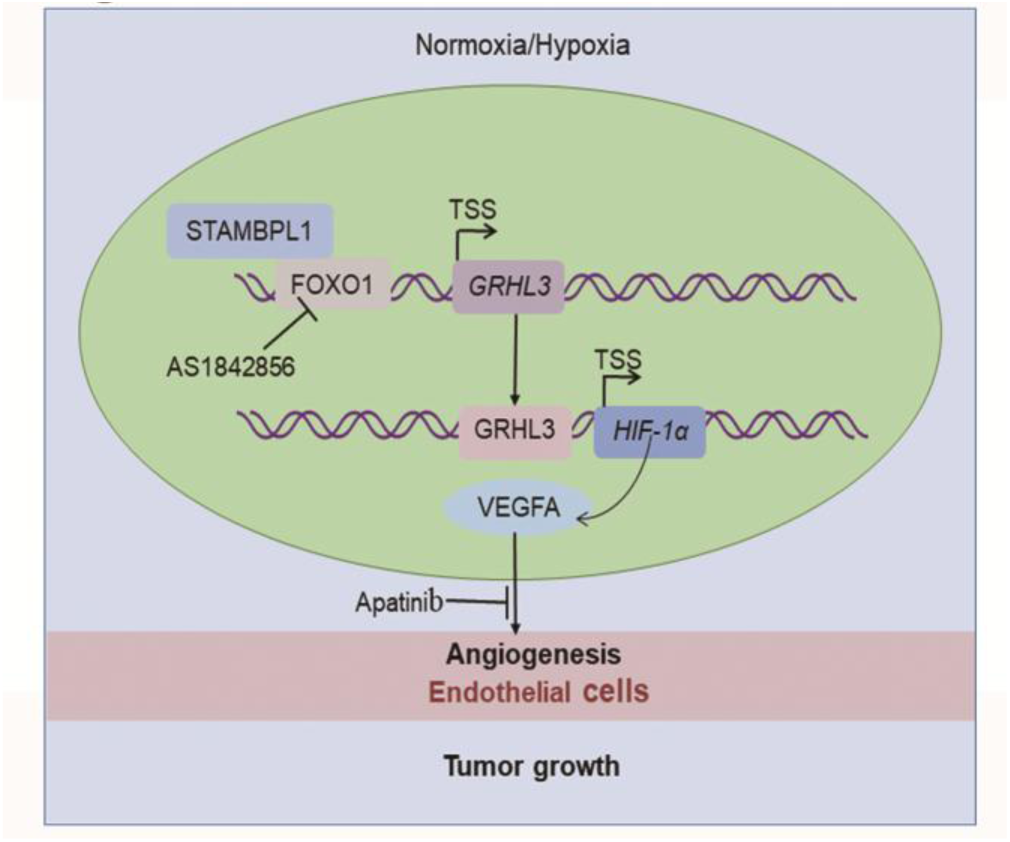
The working model of this study. STAMBPL1 interacts with FOXO1 to promote TNBC angiogenesis by activating the GRHL3/HIF1α/VEGFA axis.

## Discussion

In this study, we discovered that STAMBPL1 promotes angiogenesis in TNBC by activating the HIF1α/VEGFA pathway independently of its enzymatic activity. Through RNA-seq analysis, we revealed that STAMBPL1 positively regulates the transcription of *GRHL3*. Furthermore, we demonstrated that GRHL3 binds to the promoter of the *HIF1A* gene, thereby increasing its transcription. Additionally, we found that STAMBPL1 interacts with FOXO1 to facilitate the transcription of GRHL3/HIF1α/VEGFA. Importantly, we confirmed that the combination of the FOXO1 inhibitor AS1842856 and the VEGFR inhibitor apatinib effectively inhibited tumor growth in nude mice. These findings suggest that both STAMBPL1 and FOXO1 may be potential therapeutic targets for inhibiting angiogenesis in TNBC. This study is the first to uncover the role of STAMBPL1 in promoting angiogenesis through the FOXO1/GRHL3/HIF1α axis. Furthermore, a novel transcription factor, GRHL3, which regulates the transcription of HIF1α was identified. These findings provide valuable insights for developing new therapeutic strategies to target TNBC.

The role of STAMBPL1 in tumors has not been fully recognized. Recent studies have reported its involvement in the epithelial‒mesenchymal transition of various cancers, and its absence has been shown to affect the mesenchymal phenotype of lung and breast cancer[15]. Our previous research demonstrated that STAMBPL1 can stabilize MKP1 through deubiquitination and that the deletion of STAMBPL1 and MKP1 increases the sensitivity of breast cancer cells to cisplatin[16], suggesting that STAMBPL1 may be a potential therapeutic target for breast cancer. However, the mechanism by which STAMBPL1 activates the transcriptional activity of FOXO1 has not been elucidated. The transcriptional activity of FOXO1 is primarily regulated by its nucleocytoplasmic shuttling process[17]. The PI3K/AKT pathway promotes the phosphorylation of FOXO1, resulting in the formation of a complex with members of the 14-3-3 family (including 14-3-3σ, 14-3-3ε, and 14-3-3ζ), which facilitates its export from the nucleus and inhibits its transcriptional activity[18, 19]. It’s reported that TDAG51 prevents the binding of 14-3-3ζ to FOXO1 in the nucleus by interacting with FOXO1, thereby enhancing its transcriptional activity through increased accumulation within the nucleus[20]. Our results indicate that the overexpression of STAMBPL1 and STAMBPL1-E292A did not affect the protein levels of FOXO1 (Fig.7E and Fig.S5E), but STAMBPL1 co-localizes with FOXO1 in the nucleus (Fig.7M) and interacts with it (Fig.7N and Fig.S5I-J). This suggests that STAMBPL1 enhances the transcriptional activity of FOXO1 on GRHL3 by interacting with nuclear FOXO1. So, it will be important to develop a *Stambpl1* knockout mouse model to investigate the exact role of STAMBPL1 in TNBC. Additionally, we need to develop HIF1α and FOXO1 antibodies suitable for immunohistochemistry to detect their expression in TNBC clinical samples.

Several studies have reported that FOXO1 inhibits tumor angiogenesis[21–25]. Studies have shown that M2 macrophage-derived exosomal miR-942 promotes the migration and invasion of lung adenocarcinoma cells and facilitates angiogenesis by binding to FOXO1 to alleviate the inhibition of β-catenin, in which the upregulation of FOXO1 induces a decrease in cell invasion and angiogenesis *in vitro*[21]. FOXO1 inhibits gastric cancer growth and angiogenesis under hypoxic conditions via inactivation of the HIF1α-VEGF pathway, possibly in association with SIRT1[22]. Cancer-associated fibroblast (CAF)-derived extracellular vesicles (EVs) deliver miR-135b-5p into colorectal adenocarcinoma cells to downregulate FOXO1 and promote HUVEC proliferation, migration, and angiogenesis[23]. Colorectal cancer cell-derived exosomes overexpressing miR-183-5p promote the proliferation, migration and tube formation of HMEC-1 (human microvascular endothelial cells) cells through the inhibition of FOXO1[24]. Bladder cancer cell-derived exosomal miR-1247-3p facilitates angiogenesis by inhibiting FOXO1 expression[25]. However, the role of FOXO1 in breast cancer angiogenesis has not been studied, and our study revealed that FOXO1 can promote the expression of HIF1α and VEGFA, suggesting that it may play a role in promoting angiogenesis in breast cancer.

Studies have demonstrated that the AKT-FOXO1 signaling pathway regulates the expression of GRHL3. To investigate this, the researchers utilized a previously published ChIP-seq dataset for FOXO1 from human endometrial stromal cells. They reported that FOXO1 occupied BRD4-bound enhancers near the *GRHL3* gene. The authors subsequently confirmed that the removal of EGF and insulin from the growth medium significantly increased the expression of GRHL3, but the administration of the FOXO1 inhibitor AS1842856 partially blocked the induced expression of GRHL3[14]. These results suggest that FOXO1 plays a regulatory role upstream of GRHL3, and our study also confirmed that FOXO1 promotes the transcription of *GRHL3* by binding to its promoter.

GRHL3 is a highly conserved epidermal-specific developmental transcription factor that has recently gained attention in the field of cancer research[26]. To date, only a few genes regulated by GRHL3 have been identified. Studies have shown that GRHL3 levels are significantly reduced in human skin and head and neck squamous cell carcinomas and suggest that GRHL3 is a key tumor suppressor pathway in squamous cell carcinomas[27]. GRHL3 was also found to be induced by TNF-α in the mammary carcinoma cell line MCF-7 and was identified as a TNFα-induced endothelial cell migration factor with promigratory activity as high as that of VEGF[28]. However, the specific role of GRHL3 in breast cancer is unclear. Studies have confirmed that GRHL3 strongly stimulates endothelial cell migration, which is consistent with an angiogenic, protumorigenic function[28]. In our study, we demonstrated that GRHL3 promotes TNBC angiogenesis. HIF1α, a known regulator of angiogenesis, is regulated by growth factors[29, 30], cytokines[31] and mitogens[32, 33]. The EGFR/Akt pathway is a known positive regulator of HIF1α, potentially through mTOR[29] or independent of it[34, 35]. While the mechanisms regulating HIF1α protein expression have been extensively studied, those modulating HIF1α transcriptional activity remain unclear. Our study reveals for the first time that GRHL3 promotes transcriptional activity by binding to the promoter of the *HIF1α* gene.

In this study, we utilized two triple-negative breast cancer cell lines, HCC1806 and HCC1937, along with human primary umbilical vein endothelial cells (HUVECs) and a nude mouse breast orthotopic transplantation tumor model to investigate the regulatory mechanism by which STAMBPL1 activates the GRHL3/HIF1α/VEGFA signaling pathway through its interaction with FOXO1, thereby promoting angiogenesis in TNBC. The results of this study have certain limitations regarding their applicability to human TNBC biology. Furthermore, in addition to the HIF1α/VEGFA signaling pathway emphasized in this study, tumor cells can continuously release or upregulate various pro-angiogenic factors, such as Angiopoietin and FGF, which activate endothelial cells, pericytes (PCs), cancer-associated fibroblasts (CAFs), endothelial progenitor cells (EPCs), and immune cells (ICs). This leads to capillary dilation, basement membrane disruption, extracellular matrix remodeling, pericyte detachment, and endothelial cell differentiation, thereby sustaining a highly active state of angiogenesis[36]. It is important to collect clinical TNBC tissue samples in the future to analyze the expression of the STAMBPL1/FOXO1/GRHL3/HIF1α/VEGFA signaling axis. Furthermore, patient-derived organoid and xenograft models are useful to elucidate the regulatory relationship of this axis in TNBC angiogenesis.

### Conclusions

In summary, STAMBPL1 promoted TNBC angiogenesis by activating the GRHL3/HIF1α/VEGFA pathway via interacting with FOXO1 in a non-enzymatic manner. These findings highlight the significant role of STAMBPL1 in TNBC angiogenesis and suggest that targeting the STAMBL1/FOXO1/GRHL3/HIF1α/VEGFA axis could be a potential therapeutic strategy to inhibit angiogenesis in TNBC.

## List of abbreviations

TNBC: triple-negative breast cancer
ER: estrogen receptor
PR: progesterone receptor
HER2: human epidermal growth factor receptor 2
EGFR: epidermal growth factor receptor
PARP: poly (ADP-ribose) polymerase
VEGFA: vascular endothelial growth factor A
HUVECs: primary human umbilical vein endothelial cells
CM: conditioned medium

## Supplementary information

Supplementary Data (Supplementary Material 1) are available at eLife Online.

## Acknowledgements

We sincerely thank the team members for their dedication to this study.

## Authors’ contributions

Ceshi Chen, Huifeng Zhang, Wenmin Cao and Huichun Liang: conceptualization; Huan Fang, Huichun Liang and Ceshi Chen: data curation; Huan Fang and Huichun Liang: formal analysis, methodology and writing-original draft with help from Chuanyu Yang, Dewei Jiang and Qianmei Luo; Ceshi Chen, Huichun Liang, Huifeng Zhang, and Wenmin Cao: funding acquisition, project administration, validation, supervision and writing-review & editing.

## Declarations

### Ethics approval and consent to participate

All experimental procedures involving animals were performed in accordance to the Animal Welfare Law of China and the Guidelines for Animal Experiment of Kunming Institute of Zoology, Chinese Academy of Sciences.

### Availability of data and materials

The datasets used and/or analyzed during the current study are available from the corresponding author on reasonable request.

## Additional material

### Supplementary Material 1

**Figure S1.**
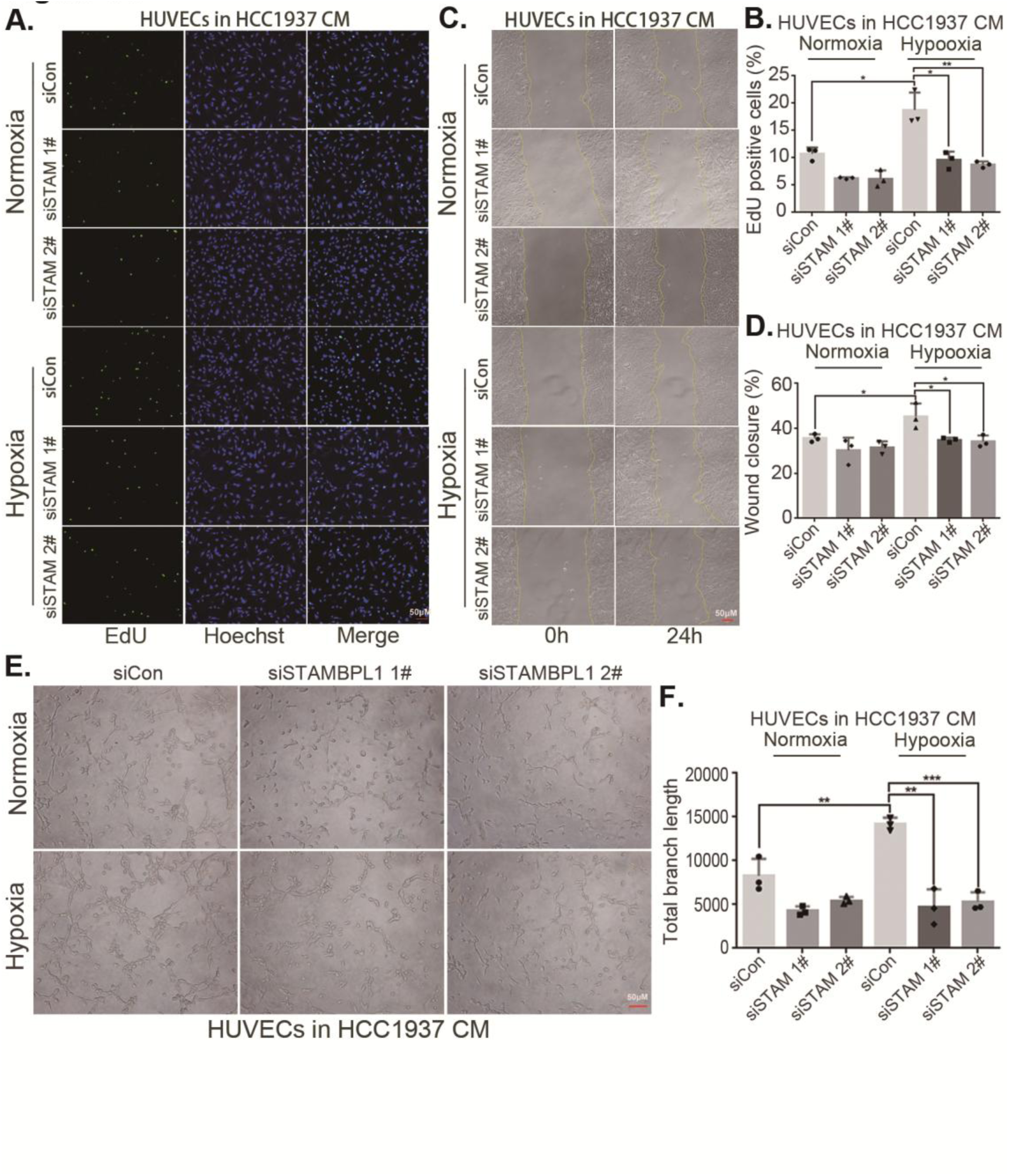
STAMBPL1 in TNBC cells enhances the activity of HUVECs. **(A)** EDU assay was performed to detect the effect of HCC1937 CM with STAMBPL1 knockdown under normoxia or hypoxia on the proliferation of HUVECs. **(B)** Statistical analysis of the EdU assay results. **(C)** Wound healing assay was performed to detect the effect of HCC1937 CM with STAMBPL1 knockdown under normoxia or hypoxia on the migration of HUVECs. **(D)** Statistical analysis of the wound healing assay results. **(E)** Tube formation assay was performed to detect the effect of HCC1937 CM with STAMBPL1 knockdown under normoxia or hypoxia on the tube formation of HUVECs. **(F)** Statistical analysis of the tube formation assay results. *: *P*<0.05, **: *P*<0.01, ***: *P*<0.001, *t-*test.

**Figure S2.**
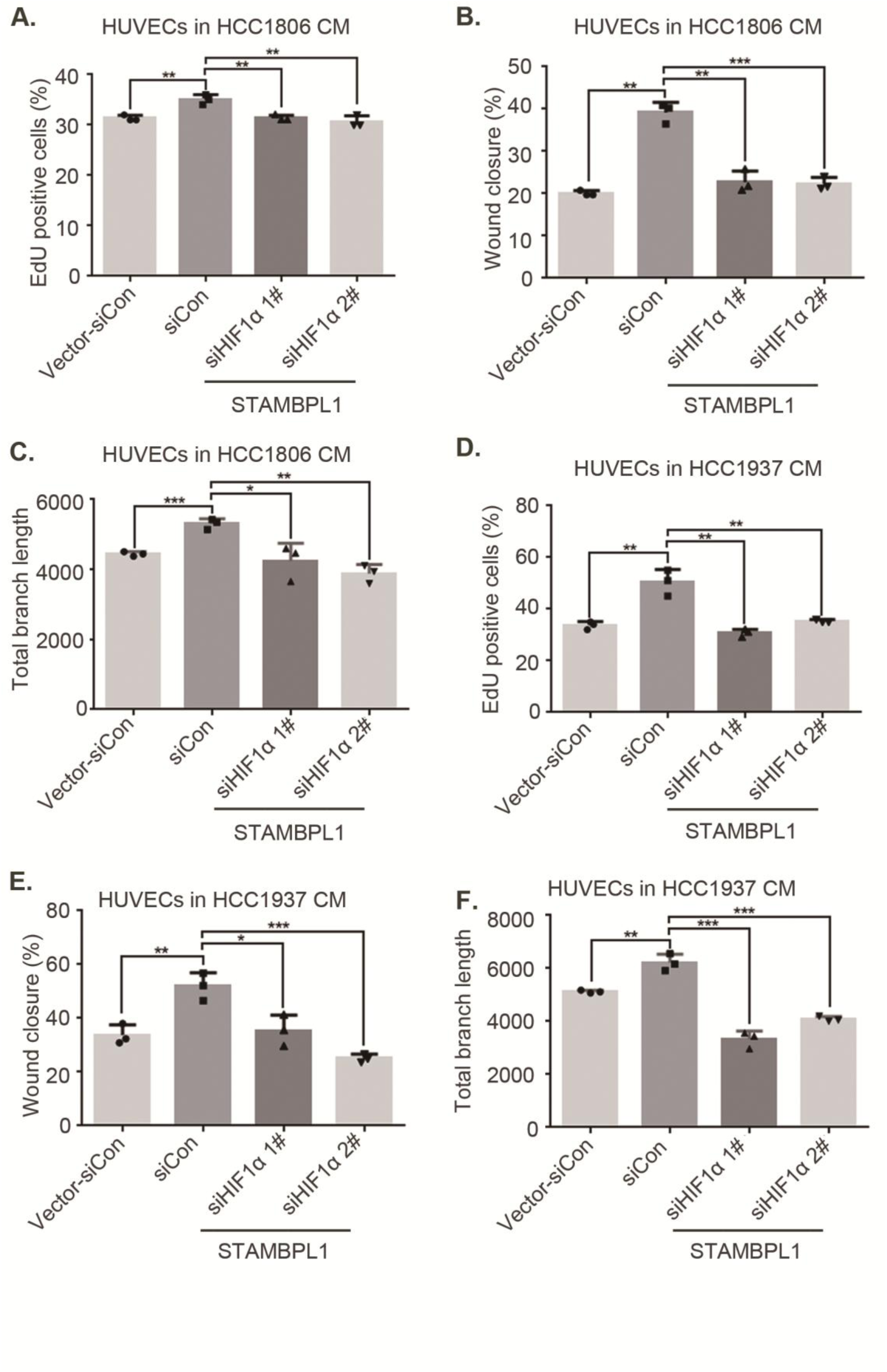
STAMBPL1 in TNBC cells enhances the activity of HUVECs. (**A and D**) HCC1806 and HCC1937 cells were stably overexpressing STAMBPL1, and HIF1α was knocked down using siRNA. The conditioned medium was then collected and used to treat HUVECs. EdU assay was performed to detect the effect of CM with HIF1α knockdown on the proliferation of HUVECs as STAMBPL1 overexpression. Statistical analysis of the EdU assay results was performed. **(B and E)** Wound healing assay was performed to detect the effect of CM with HIF1α knockdown on the migration of HUVECs as STAMBPL1 overexpression. Statistical analysis of the wound healing assay results was performed. **(C and F)** Tube formation assay was performed to detect the effect of CM with HIF1α knockdown on the tube formation of HUVECs as STAMBPL1 overexpression. Statistical analysis of the tube formation assay results was performed. *: *P*<0.05, **: *P*<0.01, ***: *P*<0.001, *t*-test.

**Figure S3.**
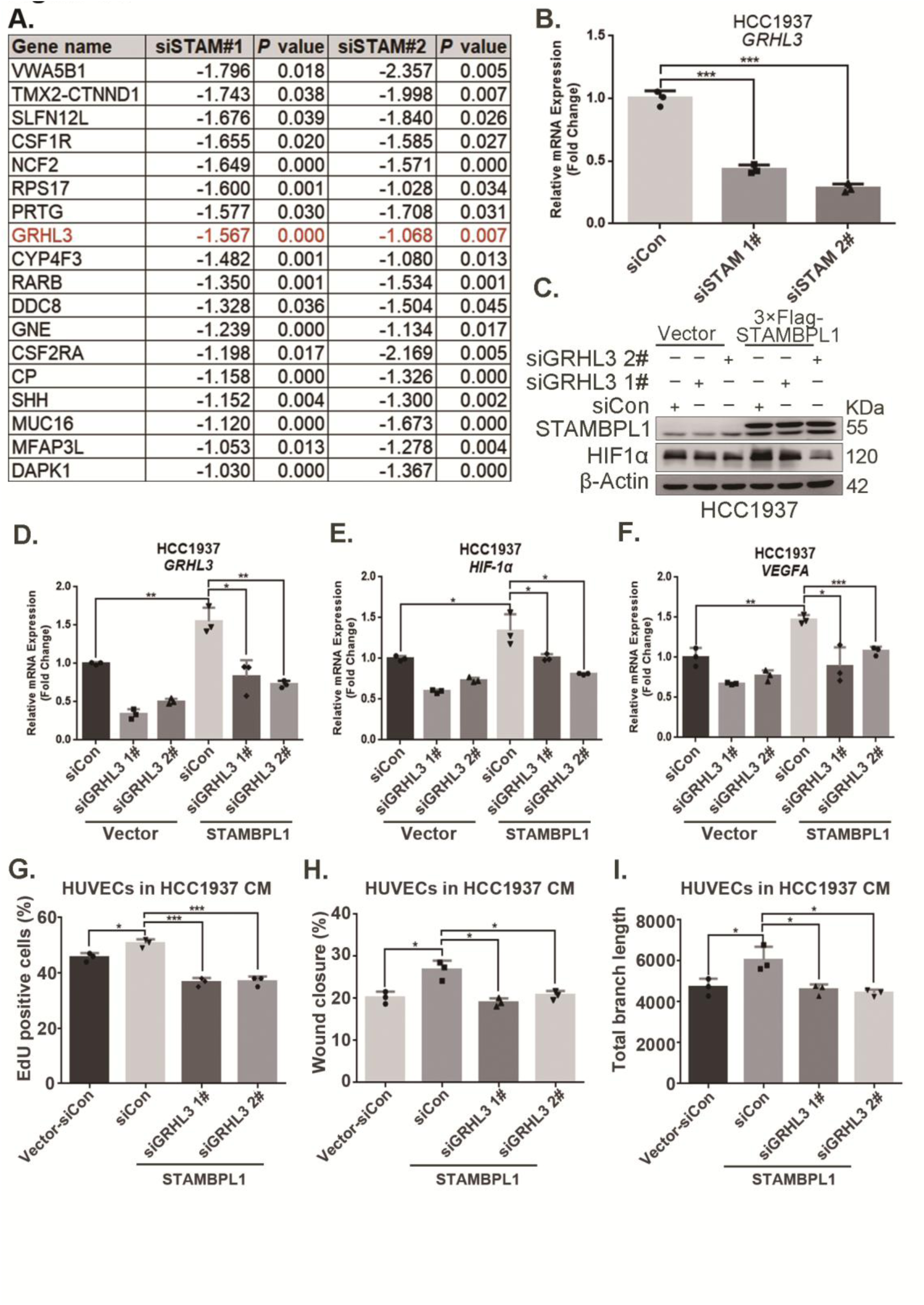
STAMBPL1 promotes *HIF1A* transcription via upregulating GRHL3. **(A)** RNA-seq result: the list of 18 genes downregulated after STAMBPL1 knockdown in HCC1806 cells. **(B)** RT-qPCR experiments showed that knockdown of STAMBPL1 inhibited GRHL3 mRNA levels in HCC1937 cells. **(C)** In HCC1937 cells stably overexpressing STAMBPL1, GRHL3 was knocked down using siRNA, and then the protein was collected. Western blotting was performed to detect the effect of GRHL3 knockdown on the expression of HIF1α as STAMBPL1overexpression. **(D-F)** RT-qPCR experiments were performed to detect the knockdown efficiency of GRHL3 and the effect of GRHL3 knockdown on the mRNA levels of HIF1α and its downstream target VEGFA when STAMBPL1 was overexpressing. **(G)** Statistical analysis of EdU assay. **(H)** Statistical analysis of wound healing assay. **(I)** Statistical analysis of tube formation assay. *: *P*<0.05, **: *P*<0.01, ***: *P*<0.001, *t*-test.

**Figure S4.**
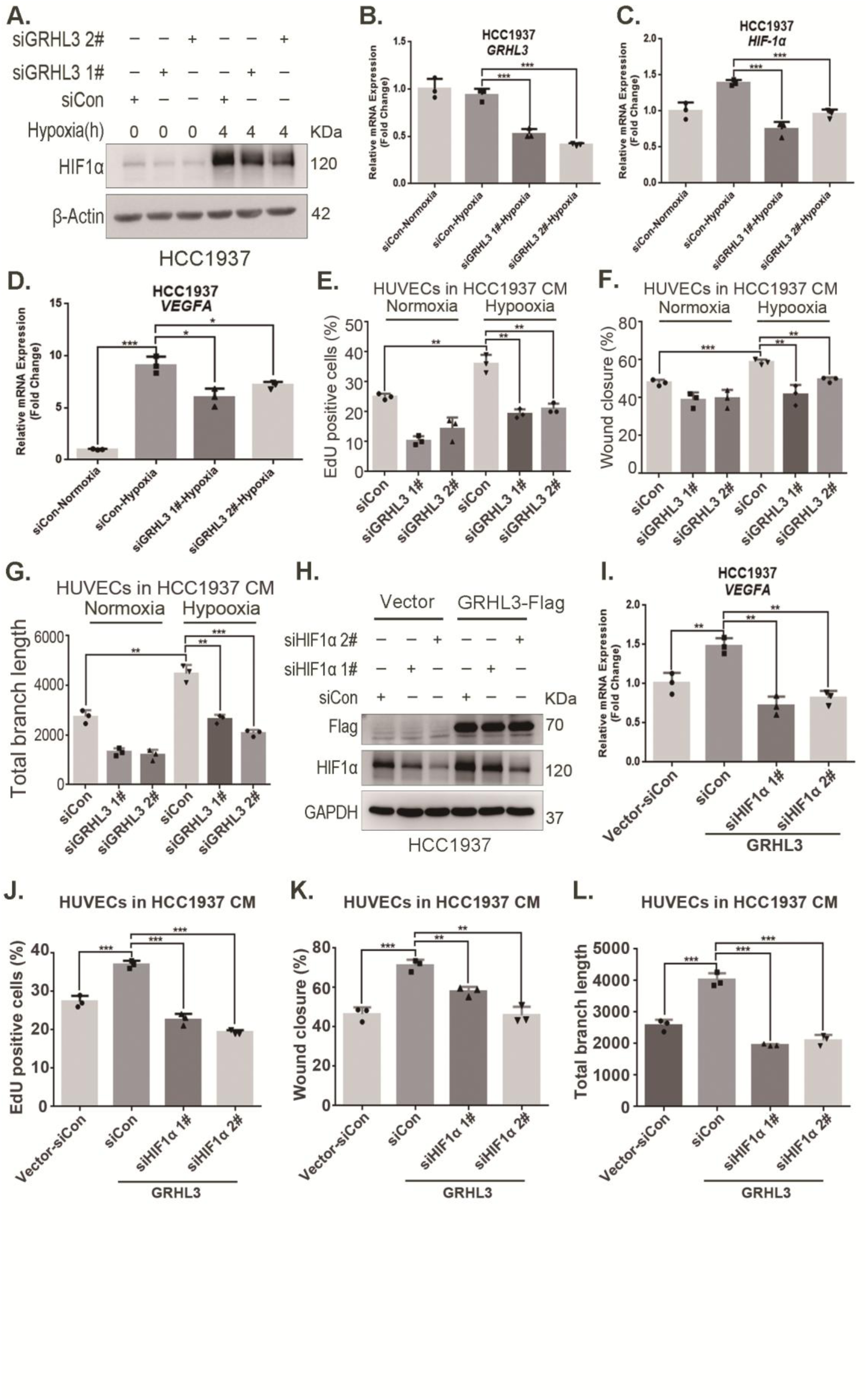
GRHL3 enhances *HIF1A* transcription by binding to its promoter. **(A)** In HCC1937 cells, knockdown of GRHL3 using siRNA and treated with hypoxia for 4 hours. Western blotting was performed to detect the effect of GRHL3 knockdown on the protein level of HIF1α. **(B-D)** RT-qPCR experiments were performed to detect the knockdown efficiency of GRHL3 and the effect of GRHL3 knockdown on the mRNA levels of HIF1α and its downstream target VEGFA. **(E)** In HCC1937 cells, GRHL3 was knocked down by siRNA and then treated with hypoxia for 24 hours, and then the conditioned medium was collected to treat HUVECs, EdU assay was performed to detect the effect of the CM with GRHL3 knockdown on the proliferation of HUVECs. **(F)** Wound healing assay was performed to detect the effect of the CM with GRHL3 knockdown on the migration of HUVECs. **(G)** The tube formation assay was performed to detect the effect of the CM with GRHL3 knockdown on the tube formation of HUVECs. **(H)** In HCC1937 cells stably overexpressing GRHL3, HIF1α was knocked down using siRNA, and then the protein was collected. Western blotting was performed to detect the protein levels of GRHL3 and HIF1α. **(I)** RT-qPCR experiment was performed to detect the effect of HIF1α knockdown on the mRNA expression of VEGFA as GRHL3 overexpression. **(J)** Statistical analysis of EdU assay results. **(K)** Statistical analysis of wound healing assay results. **(L)** Statistical analysis of tube formation assay results. *: *P*<0.05, **: *P*<0.01, ***: *P*<0.001, *t*-test.

**Figure S5.**
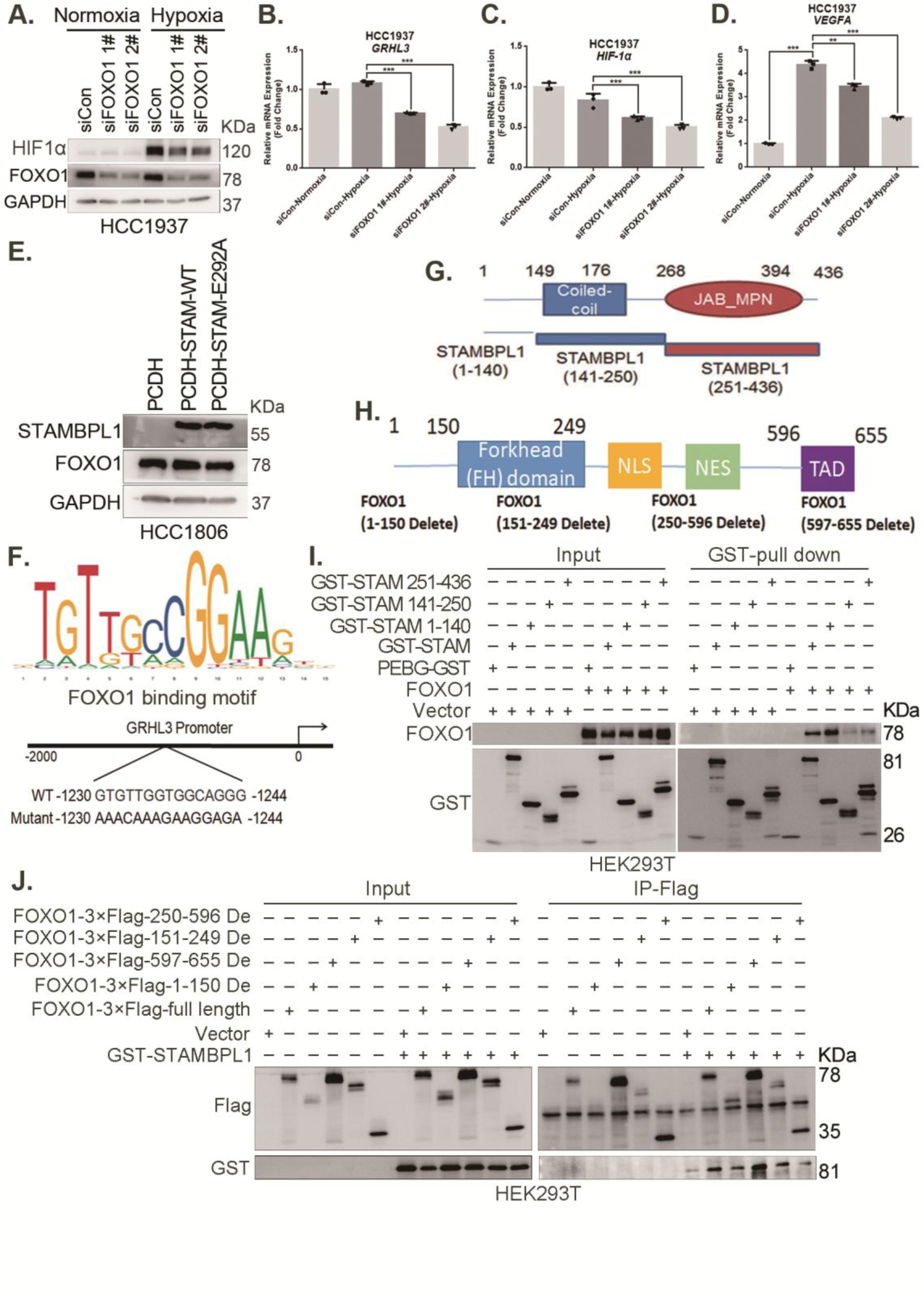
STAMBPL1 mediates *GRHL3* transcription by interacting with FOXO1. **(A)** Western blotting was performed to detect the effect of FOXO1 knockdown on the protein level of HIF1α in HCC1937 cells under hypoxia for 4 hours. **(B-D)** RT-qPCR experiments were performed to detect the effect of FOXO1 knockdown on the mRNA levels of GRHL3/HIF1α/VEGFA in HCC1937 cells under hypoxia treatment for 4 hours. **(E)** Western blot detected the FOXO1 expression when STAMBPL1 and STAMPBL1-E292A were overexpressed. **(F)** The JASPAR website predicts the possible binding sequence of transcription factor FOXO1 to the GRHL3 promoter and its mutation pattern map. **(G)** The truncated structure diagram of STAMBPL1. **(H)** The truncated structure diagram of FOXO1. **(I)** The full-length and truncated plasmids of pEBG-STAMBPL1-GST and pCDH-FOXO1-no tag were transfected into HEK293T cells, and the proteins were collected for GST-pull down assay at 48 hours after transfection. **(J)** The full-length and truncated plasmids of pCDH-FOXO1-3×Flag and pEBG-STAMBPL1-GST were transfected into HEK293T cells, and the proteins were collected for IP-Flag assay at 48 hours after transfection. *: *P*<0.05, **: *P*<0.01, ***: *P*<0.001, *t*-test.

**Figure S6.**
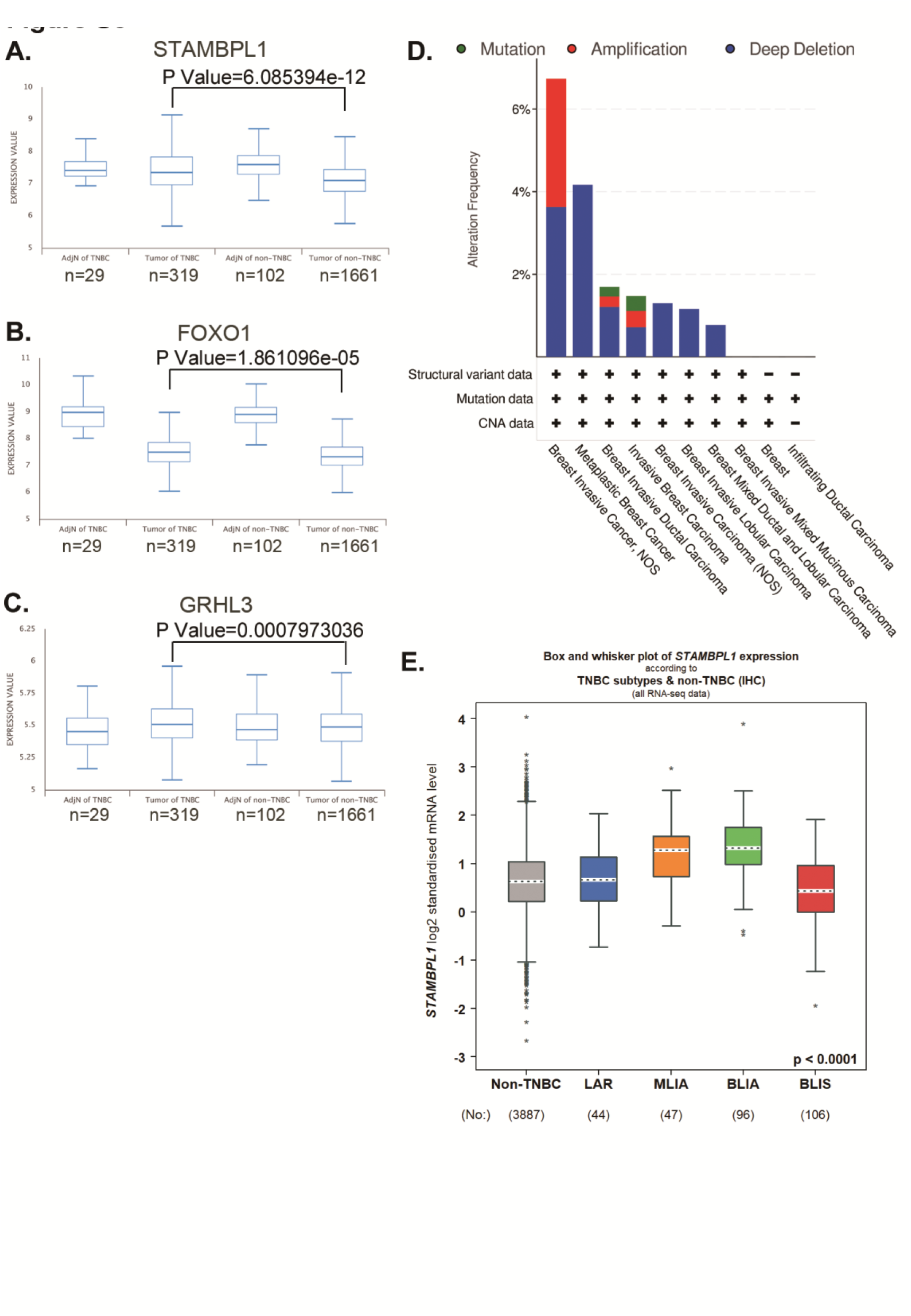
STAMBPL1, FOXO1, and GRHL3 is highly expressed in TNBC. (**A-C**) The expression of STAMBPL1, FOXO1, and GRHL3 in adjacent normal of TNBC (n=29), TNBC (n=319), adjacent normal of non-TNBC (n=102), and non-TNBC (n=1661) by analyzing the Metabric database in BCIP. **(D)** The mutation, amplification, and deep deletion of STAMBPL1 in breast cancer were analyzed by using cBioportal online database. **(E)** The expression of STAMBPL1 in non-TNBC and subtypes of TNBC by using bc-GenExMiner online database.

