## Supplementary Material 1 for "STAMBPL1 activates the GRHL3/HIF1A/VEGFA axis through interaction with FOXO1 to promote angiogenesis in triple-negative breast cancer"

### Figure S1

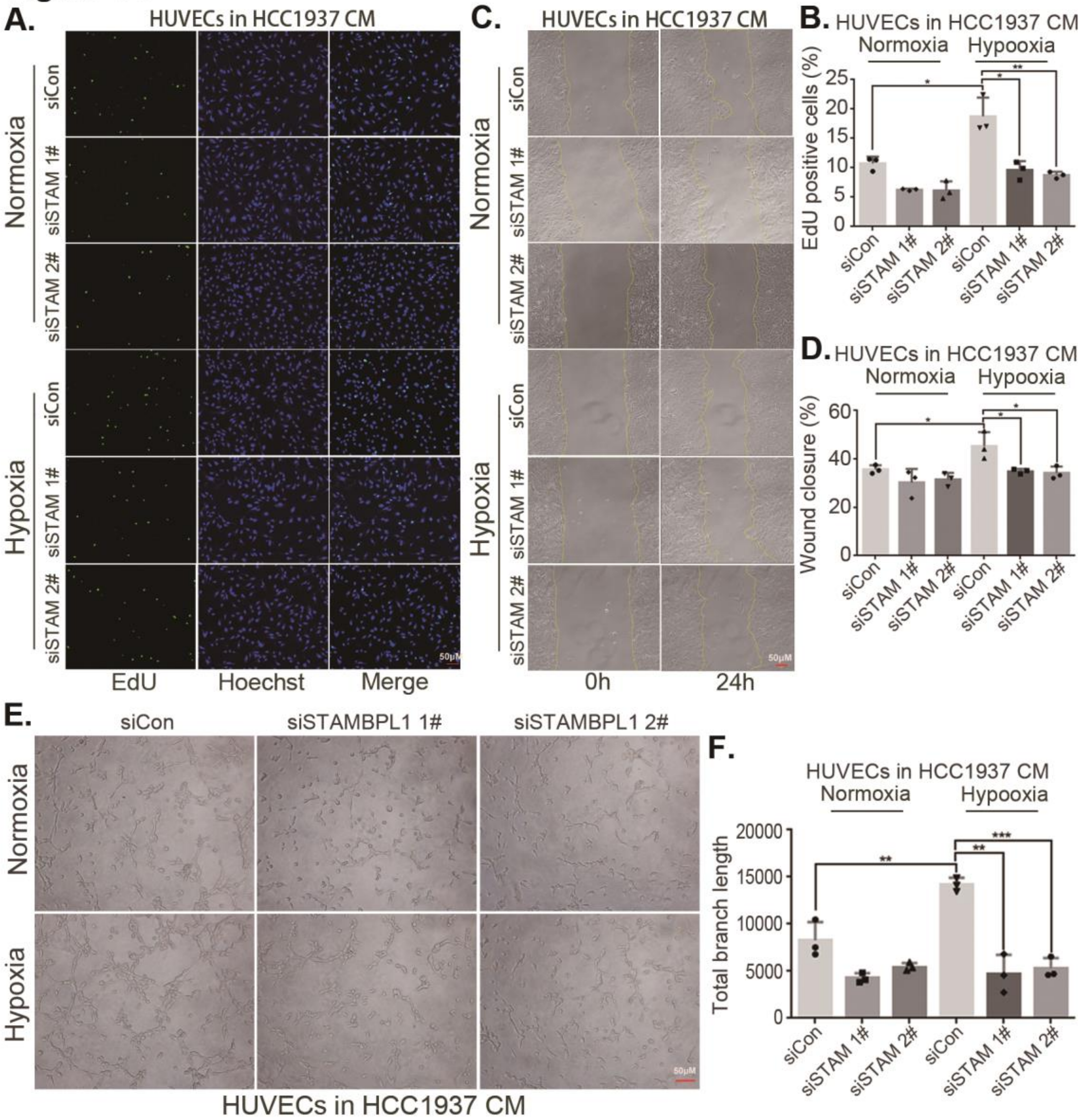
A.
HUVECs in HCC1937 CM
Normoxia
Hypoxia
siCon
siSTAM 1#
siSTAM 2#
EdU
Hoechst
Merge
50μM
C.
HUVECs in HCC1937 CM
0h
24h
B. HUVECs in HCC1937 CM
Normoxia
Hypooxia
EdU positive cells (%)
D. HUVECs in HCC1937 CM
Wound closure (%)
E.
siCon
siSTAMBPL1 1#
siSTAMBPL1 2#
HUVECs in HCC1937 CM
F.
HUVECs in HCC1937 CM
Total branch length

**Figure S2**

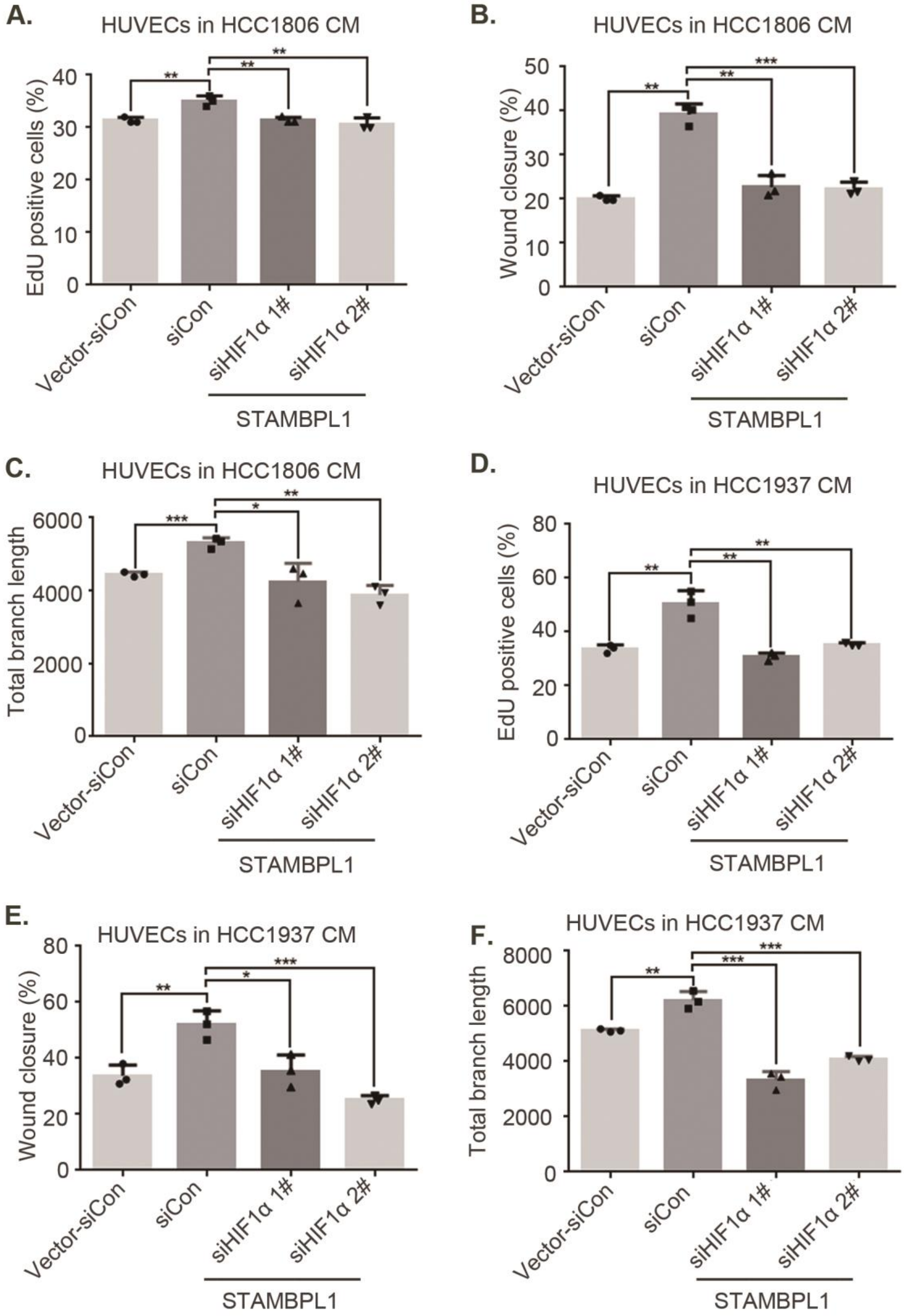
A.
HUVECs in HCC1806 CM
EdU positive cells (%)
Vector-siCon
siCon
siHIF1α 1#
siHIF1α 2#
STAMBPL1
B.
HUVECs in HCC1806 CM
Wound closure (%)
C.
HUVECs in HCC1806 CM
Total branch length
D.
HUVECs in HCC1937 CM
EdU positive cells (%)
E.
HUVECs in HCC1937 CM
Wound closure (%)
F.
HUVECs in HCC1937 CM
Total branch length

#### Figure S3

**A.**

| Gene name | siSTAM#1 | *P* value | siSTAM#2 | *P* value |
|---|---|---|---|---|
| VWA5B1 | -1.796 | 0.018 | -2.357 | 0.005 |
| TMX2-CTNND1 | -1.743 | 0.038 | -1.998 | 0.007 |
| SLFN12L | -1.676 | 0.039 | -1.840 | 0.026 |
| CSF1R | -1.655 | 0.020 | -1.585 | 0.027 |
| NCF2 | -1.649 | 0.000 | -1.571 | 0.000 |
| RPS17 | -1.600 | 0.001 | -1.028 | 0.034 |
| PRTG | -1.577 | 0.030 | -1.708 | 0.031 |
| GRHL3 | -1.567 | 0.000 | -1.068 | 0.007 |
| CYP4F3 | -1.482 | 0.001 | -1.080 | 0.013 |
| RARB | -1.350 | 0.001 | -1.534 | 0.001 |
| DDC8 | -1.328 | 0.036 | -1.504 | 0.045 |
| GNE | -1.239 | 0.000 | -1.134 | 0.017 |
| CSF2RA | -1.198 | 0.017 | -2.169 | 0.005 |
| CP | -1.158 | 0.000 | -1.326 | 0.000 |
| SHH | -1.152 | 0.004 | -1.300 | 0.002 |
| MUC16 | -1.120 | 0.000 | -1.673 | 0.000 |
| MFAP3L | -1.053 | 0.013 | -1.278 | 0.004 |
| DAPK1 | -1.030 | 0.000 | -1.367 | 0.000 |

**B.**

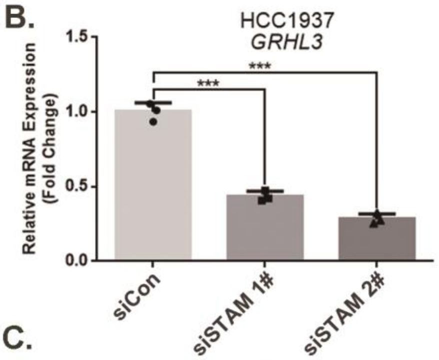


**C.**

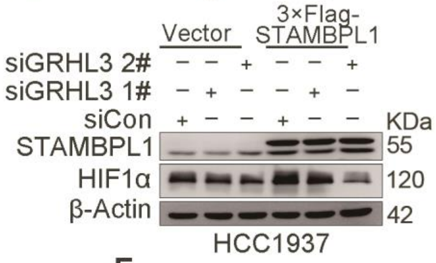


**D.** **E.** **F.**

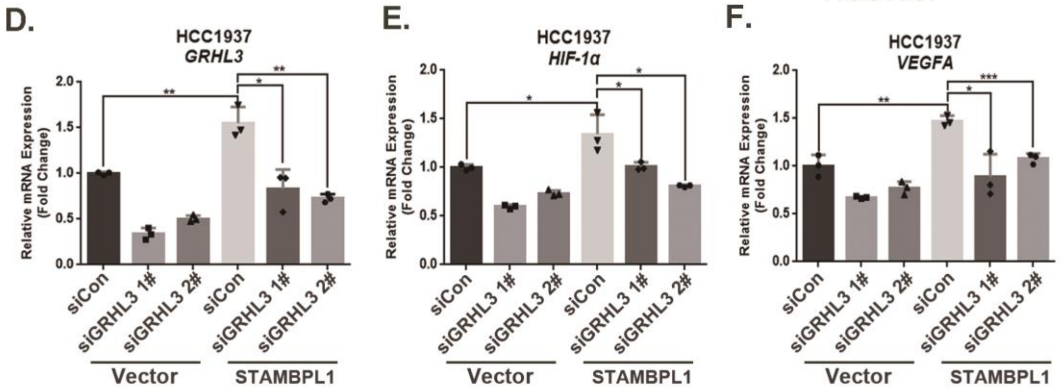


**G.** **H.** **I.**

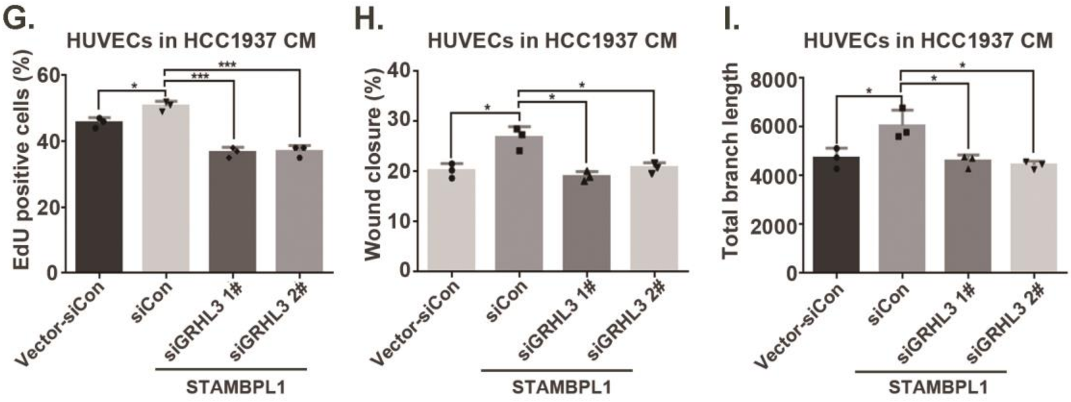

#### Figure S4

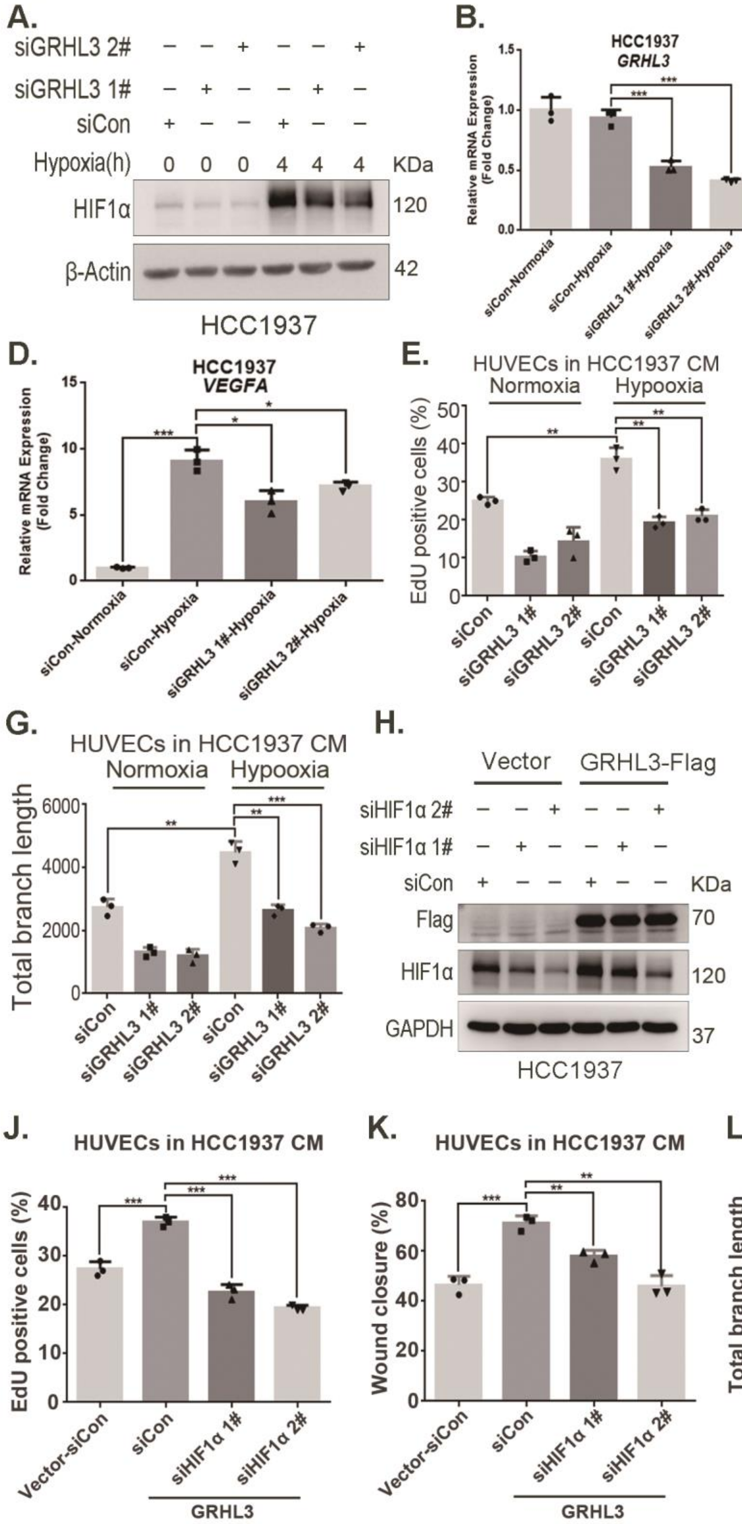

A.
siGRHL3 2#
siGRHL3 1#
siCon
Hypoxia(h)
KDa
HIF1α
β-Actin
HCC1937
B.
HCC1937
GRHL3
Relative mRNA Expression (Fold Change)
siCon-Normoxia
siCon-Hypoxia
siGRHL3 1#-Hypoxia
siGRHL3 2#-Hypoxia
C.
HCC1937
HIF-1α
D.
HCC1937
VEGFA
E.
HUVECs in HCC1937 CM
Normoxia
Hypooxia
EdU positive cells (%)
siCon
siGRHL3 1#
siGRHL3 2#
F.
HUVECs in HCC1937 CM
Wound closure (%)
G.
HUVECs in HCC1937 CM
Total branch length
H.
Vector
GRHL3-Flag
siHIF1α 2#
siHIF1α 1#
Flag
GAPDH
I.
HCC1937
VEGFA
Vector-siCon
GRHL3
J.
HUVECs in HCC1937 CM
K.
HUVECs in HCC1937 CM
L.
HUVECs in HCC1937 CM

#### Figure S5

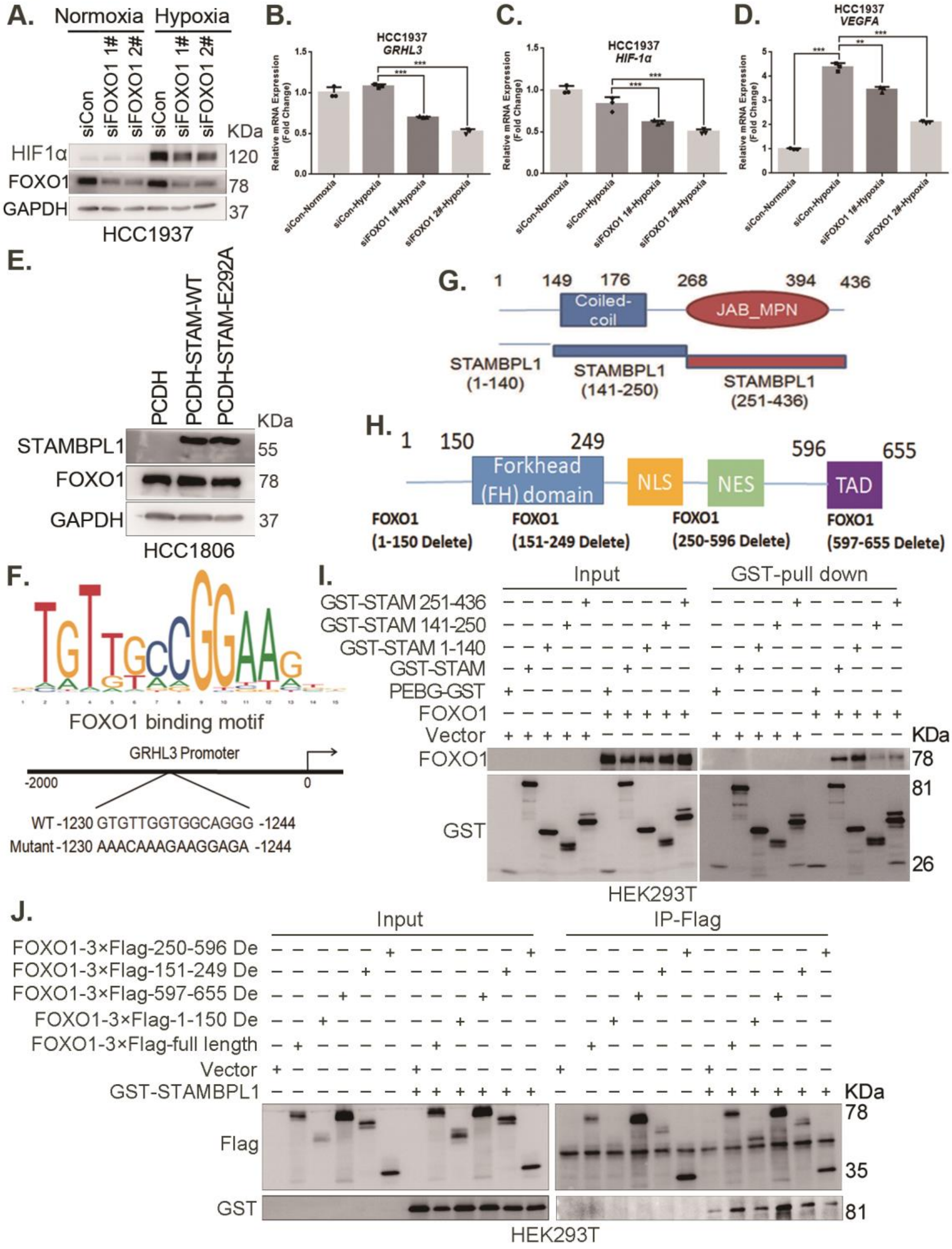

A.
Normoxia
Hypoxia
siCon
siFOXO1 1#
siFOXO1 2#
KDa
HIF1α
FOXO1
GAPDH
120
78
37
HCC1937
B.
HCC1937
GRHL3
C.
HCC1937
HIF-1α
D.
HCC1937
VEGFA
E.
PCDH
PCDH-STAM-WT
PCDH-STAM-E292A
STAMBPL1
55
HCC1806
F.
FOXO1 binding motif
GRHL3 Promoter
-2000
0
WT -1230 GTGTTGGTGGCAGGG -1244
Mutant -1230 AAACAAAGAAGGAGA -1244
G.
Coiled-coil
JAB_MPN
STAMBPL1 (1-140)
STAMBPL1 (141-250)
STAMBPL1 (251-436)
H.
Forkhead (FH) domain
NLS
NES
TAD
FOXO1 (1-150 Delete)
FOXO1 (151-249 Delete)
FOXO1 (250-596 Delete)
FOXO1 (597-655 Delete)
I.
Input
GST-pull down
GST-STAM 251-436
GST-STAM 141-250
GST-STAM 1-140
GST-STAM
PEBG-GST
FOXO1
Vector
GST
HEK293T
J.
Input
IP-Flag
FOXO1-3×Flag-250-596 De
FOXO1-3×Flag-151-249 De
FOXO1-3×Flag-597-655 De
FOXO1-3×Flag-1-150 De
FOXO1-3×Flag-full length
Vector
GST-STAMBPL1
Flag
GST
HEK293T

#### Figure S6

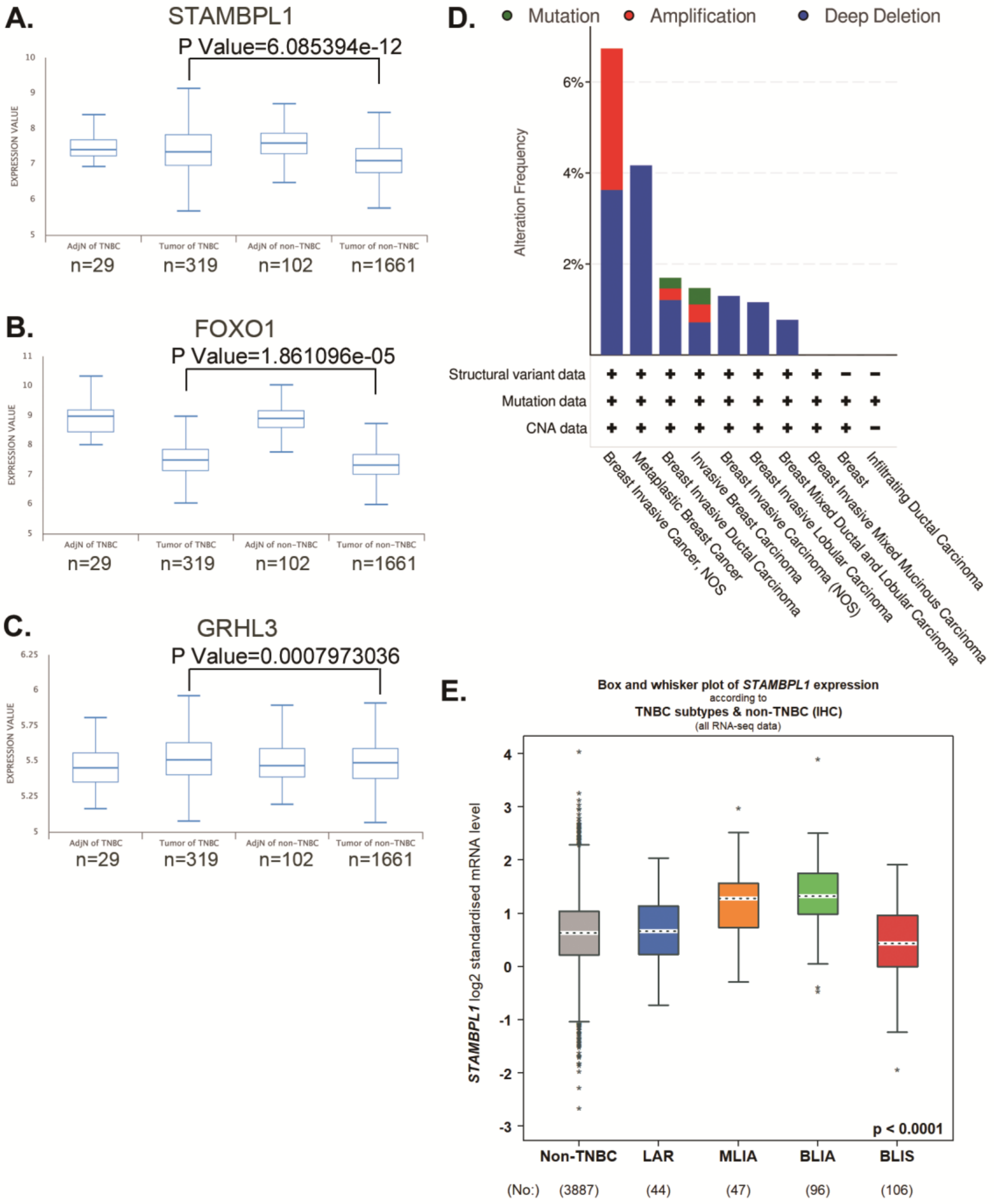
A.
STAMBPL1
P Value=6.085394e-12
EXPRESSION VALUE
AdjN of TNBC
Tumor of TNBC
AdjN of non-TNBC
Tumor of non-TNBC
n=29
n=319
n=102
n=1661
B.
FOXO1
P Value=1.861096e-05
EXPRESSION VALUE
AdjN of TNBC
Tumor of TNBC
AdjN of non-TNBC
Tumor of non-TNBC
n=29
n=319
n=102
n=1661
C.
GRHL3
P Value=0.0007973036
EXPRESSION VALUE
AdjN of TNBC
Tumor of TNBC
AdjN of non-TNBC
Tumor of non-TNBC
n=29
n=319
n=102
n=1661
D.
Mutation
Amplification
Deep Deletion
Alteration Frequency
6%
4%
2%
Structural variant data
Mutation data
CNA data
Breast Invasive Cancer, NOS
Metaplastic Breast Cancer
Breast Invasive Ductal Carcinoma
Invasive Breast Carcinoma
Breast Invasive Carcinoma (NOS)
Breast Invasive Lobular Carcinoma
Breast Mixed Ductal and Lobular Carcinoma
Breast Invasive Mixed Mucinous Carcinoma
Breast
Infiltrating Ductal Carcinoma
E.
Box and whisker plot of STAMBPL1 expression
according to
TNBC subtypes & non-TNBC (IHC)
(all RNA-seq data)
STAMBPL1 log2 standardised mRNA level
p < 0.0001
Non-TNBC
LAR
MLIA
BLIA
BLIS
(No:)
(3887)
(44)
(47)
(96)
(106)
